# A haplotype-resolved T2T genome assembly of Indigofera pseudotinctoria reveals the genetic basis of flavonoid biosynthesis in Chinese Indigo

**DOI:** 10.64898/2025.12.05.692270

**Authors:** Jinghan Peng, Junming Zhao, Jie Zhou, Yi xiong, Yuandong Xu, Huizhen Ma, Jing Chen, Qifan Ran, Wei He, Xiao Ma, Yan Fan

## Abstract

*Indigofera pseudotinctoria,* commonly known as Chinese Indigo, is a multifunctional leguminous shrub widely distributed in East Asia, valued for its medicinal, ecological, and forage importance, and naturally adapted to drought and nutrient-poor soils. However, the absence of genomic resources has hindered insights into the genetic basis of its bioactive metabolites and adaptive traits. Here, we present a haplotype-resolved, telomere-to-telomere (T2T) reference genome of *I. pseudotinctoria*, which represents the first complete genome within the genus Indigofera. By integrating PacBio HiFi sequencing, Oxford Nanopore ultra-long reads, and Hi-C chromatin conformation sequencing technologies, we successfully constructed two gap-free haplotypes assemblies (647.38 Mb and 657.52 Mb), characterized by high BUSCO completeness and QV scores, and fully captured all telomeric sequences. Comparative analysis between haplotypes revealed extensive synteny but notable structural heterozygosity but notable structural heterozygosity. Approximately 46% of structural variations overlap with genic or regulatory regions, leading to allele-specific expression divergence. Between the two haplotypes, we identified 2,607 and 2,331 haplotype-specific genes, reflecting a complementary functional specialization in defense and repair versus metabolism and growth. Integrative transcriptomic and metabolomic profiling across six tissues reconstructed the flavonoid biosynthetic network and identified MYB, bHLH, NAC, WRKY, and ERF transcription factors regulating five key pharmacologically active flavonoids (Calycosin, Butein, Sulfuretin, Chrysoeriol, and Genistin). These results collectively uncover the haplotype-specific regulatory and structural basis of Indigofera pseudotinctoria and establish a high-quality genomic framework for evolutionary, medicinal, and metabolic engineering studies in leguminous plants, and provide a framework for molecular breeding and bioactive compound discovery in medicinal legumes.

## Introduction

*Indigofera* L., the third-largest genus in the Fabaceae family, comprises ∼750 species with remarkable ecological and medicinal diversity across tropical and subtropical regions^1–3^. Among them, *Indigofera pseudotinctoria* (Chinese Indigo), a perennial diploid shrub (2n = 2x = 16) native to East Asia, stands out as a multifunctional species with profound ethnopharmacological significance^4–6^. For centuries, its roots, flowers, and pods have been integral to traditional Chinese medicine, prescribed for detoxification, anti-inflammatory, analgesic, and antioxidant therapies^7–9^. Modern pharmacology validates these uses, attributing its efficacy to a rich repertoire of bioactive flavonoids such as Calycosin, Butein, and Genistein^10,11^, which exhibit potent antioxidant, anti-inflammatory, and immunomodulatory activities^8,12^. Beyond medicine, *I. pseudotinctoria* is a high-quality forage shrub, with crude protein content rivaling alfalfa, making it invaluable for sustainable livestock husbandry, particularly in agro-pastoral ecotones^13–15^. Its drought tolerance, phytoremediation potential, and notable ornamental value further highlight its comprehensive role in ecological restoration^16,17^. However, despite these multifaceted values, the absence of a reference genome has hindered functional genetics and breeding efforts in *Indigofera*^18^.

Flavonoids constitute the predominant class of bioactive small molecules in *I. pseudotinctoria*. In legumes, the flavonoid/isoflavonoid network underpins pigmentation, signaling, and stress responses, and represents key chemotypes with antioxidant, anti-inflammatory, antimicrobial, and cytoprotective activities^18–20^. These metabolites originate from the phenylpropanoid pathway, with phenylalanine as the primary precursor. Enzymatic steps catalyzed by phenylalanine ammonia-lyase (PAL), cinnamate 4-hydroxylase (C4H), and 4-coumarate-CoA ligase (4CL) generate the essential intermediates for flavonoid biosynthesis, which are subsequently diversified through structural genes such as chalcone synthase (CHS), chalcone isomerase (CHI), flavanone 3-hydroxylase (F3H), and anthocyanidin synthase (ANS)^21,22^. Despite decades of phytochemical and pharmacological research in Indigofera^23^, the genetic architecture and regulatory logic underlying flavonoid accumulation, its variation across tissues and environments, and its links to adaptive performance remain poorly understood^8^. Furthermore, as an insect-pollinated allogamous species, *I. pseudotinctoria* exhibits extensive heterozygosity and pronounced haplotype divergence^23^. Under such genomic complexity, conventional “mosaic” genome assemblies obscure allelic and structural variants that mediate the balance between metabolic stability and allelic differentiation^24–26^. Therefore, generating a chromosome-scale, haplotype-resolved genome is essential to disentangle these layers of variation, reconstruct the shared core regulatory networks of flavonoid metabolism, and elucidate how *I. pseudotinctoria* maintains a stable supply of bioactive compounds amid a dynamic genomic landscape.

*I. pseudotinctoria* demonstrates significant heterozygosity, also offering a unique opportunity to investigate the regulatory mechanisms of alleles in outcrossing Indigofera species. A haplotype-resolved assembly enables precise identification of structural variants (SVs) and haplotype-specific genes, revealing processes such as gene duplication, subfunctionalization, and adaptive divergence at the allelic level^27–29^. Moreover, by distinguishing transcripts derived from alternative haplotypes, allele-specific expression (ASE) analysis directly reveals interactions between alleles^30,31^. Together, these insights offer a framework for understanding how genomic architecture and allelic interactions shape phenotypic robustness and adaptive potential in plants. The haplotype-resolved genome of *I. pseudotinctoria* thus provides a foundational resource for decoding allelic complexity, structural diversity, and expression bias in highly heterozygous Fabaceae species.

Here, we report the first telomere-to-telomere (T2T), haplotype-resolved genome of *I. pseudotinctoria*, constructed using PacBio HiFi, Oxford Nanopore, and Hi-C data. By integrating this genomic framework with transcriptomics and metabolomics, we pinpoint the haplotype-biased regulators and candidate genes driving the biosynthesis of five pharmacologically pivotal flavonoids. This study provides a paradigm for how T2T haplotype genomics can unlock the genetic basis of complex traits in medicinal legume plants, establishing a foundational resource for evolutionary research, functional genomics, and molecular breeding in *Indigofera* and beyond.

## Results

### The haplotype-resolved gap-free T2T genome assembly

To assemble gap-free, T2T haplotypes of *I. pseudotinctoria*, we generated high-coverage PacBio HiFi reads, Oxford Nanopore Technology (ONT) ultra-long reads, and high-throughput chromatin conformation capture (Hi-C) sequence reads (Supplementary Table1). Genome assembly was performed using an in-house pipeline (Supplementary Fig.1) as follows. Firstly, HiFiasm^32^ was used to generate a haplotype-resolved draft genome from ONT reads, HiFi reads, and Hi-C data. After removing microbial and plastid sequences, the overlapping clusters were anchored onto the 8 pseudochromosomes of the two haplotypes using Hi-C data (Supplementary Fig.2). Haplotype 1 (Hap1) contained a single gap on chromosome 8, whereas haplotype 2 (Hap2) contained three gaps, located on chromosomes 1, 5, and 8, respectively. All gaps were subsequently closed using ONT and HiFi raw reads and continuous sequences assembled with HiCanu^33^ and NextDenovo^34^.(Supplementary Fig.3 and Supplementary Table2). The final genome sizes were 647.38 Mb and 657.52 Mb for Hap1 and Hap2 (Fig. 1B and 1C, Table 1).

**Fig. 1.**
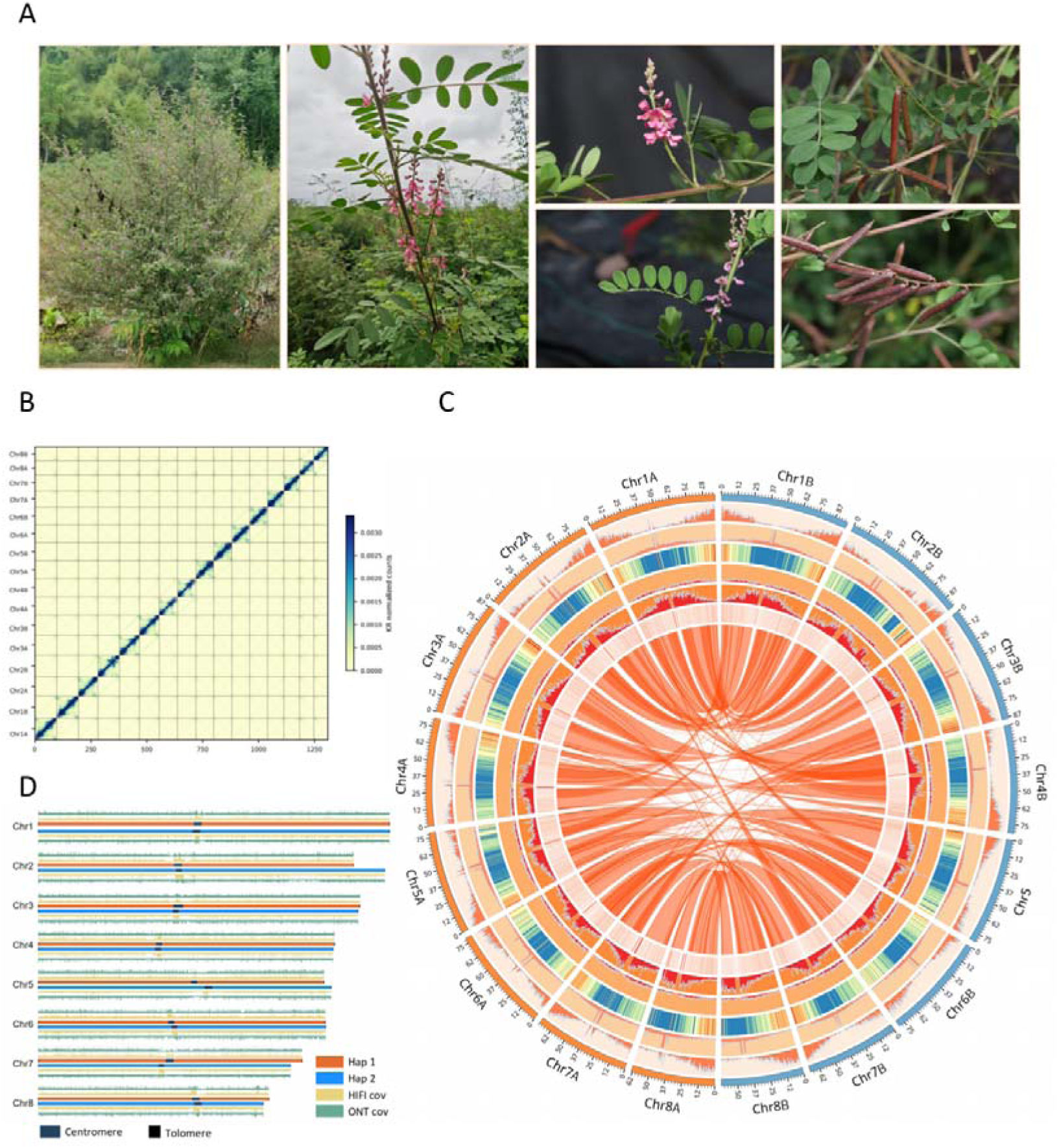
(A) The morphological characteristics of *I. pseudotinctoria*. (B) Hi-C interaction map of *I. pseudotinctoria*, with blue boxes representing individual chromosomes. The x- and y-axes indicate the ordered positions of the chromosomes of the genome. (C) Circos diagram of two haplotypes. The concentric circles from outer to inner diameters (Hap1, orange; Hap2, blue) are annotated as follows: (I) chromosome number, (II) GC content, (III) gene density, (IV) transposable element (TE) density, (V) genes collinear blocks. GC content was calculated by determining the proportion of (G + C) nucleotides within 300 kb non-overlapping windows. Gene density, TE density, SNP density, INDEL density, PAV density, and CNV density were each calculated within non-overlapping 100 kb windows. (D) Haplotype-resolved telomere-to-telomere genome assembly and sequencing coverage of *I. pseudotinctoria*. The coverage tracks represent HiFi and ONT sequencing depths.

**Table 1.** Summary of genome assembly and annotation of *I. pseudotinctoria*.

| Assembly | Hap1 | Hap2 |
| --- | --- | --- |
| Total length (Mb) | 647.38 | 657.52 |
| Largest contig (Mb) | 96.69 | 97.45 |
| Contig N50 (Mb) | 69.01 | 69.98 |
| Gap | 0 | 0 |
| Number of telomeres | 16 | 16 |
| GC content of the genome (%) | 36.93 | 36.88 |
| Quality value(QV) | 71.25 | 73.64 |
| LTR Assembly Index | 20.94 | 21.6 |
| Annotation |  |  |
| Number of predicted protein-coding genes | 27912 | 27636 |
| Masked repeat sequence length (Mb) | 449.4 | 459.29 |
| Percentage of repeat sequence sequences | 69.4% | 69.9% |
| BUSCO completeness of annotation | 98.2% | 98.5% |

Hap1 contains 27,912 protein-coding genes, along with 2,935 snoRNAs, 1,558 tRNAs, and 292 miRNAs. Hap2 contains 27,636 protein-coding genes, along with 2,724 snoRNAs, 1,558 tRNAs, and 270 miRNAs. Remarkably, rDNA clusters were detected on Chr1, Chr6, and Chr8 in both haplotypes(Supplementary Fig.4). Analysis of repetitive sequences revealed that Hap1 and Hap2 harbor 449.40 Mb and 459.29 Mb of transposable elements (TEs), respectively (Supplementary Table3), with LTRs being the most abundant TE category (Supplementary Fig.5). Following genome annotation, we evaluated the completeness and accuracy of the haplotype assemblies using multiple quantitative metrics, including contig N50, k-mer completeness, assembly Quality Value (QV) scores, and the Benchmarking Universal Single-Copy Orthologs (BUSCO) gene sets. The contig N50 values were 69.01 Mb for Hap1 and 69.98 Mb for Hap2. K-mer–based analysis indicated an overall completeness of 99.29%. The QV scores were 71.25 for Hap1 and 73.64 for Hap2, reflecting high base-level accuracy(Supplementary Table4). BUSCO analysis showed that 98.2% and 98.5% of the conserved single-copy orthologs were recovered in Hap1 and Hap2, respectively, indicating nearly complete gene space coverage. The LTR Assembly Index (LAI) values were 20.94 for Hap1 and 21.60 for Hap2, further supporting the high assembly quality for repetitive regions.

We analyzed telomeres and centromeres of the *I. pseudotinctoria* haplotype genomes. Using the telomeric repeat sequence (TTTAGGG) as a query, we identified 16 telomeres at the ends of the 8 chromosomes in Hap1 and 16 telomeres in Hap2 (Fig. 1D, Supplementary Fig. 6 and Supplementary Table 5). In summary, we successfully assembled the T2T haplotype-resolved genome and identified all telomeres, and the putative centromeric regions of the all 8 chromosomes in each haploid genome were identified (Supplementary Table 6 and 7). Centromere sequence similarity heatmaps and methylation boundary analysis supported the accuracy of centromere identification (Supplementary Fig. 7 and 8).

### Haplotype-resolved genome comparison

The haplotype-resolved genomes of *I. pseudotinctoria* exhibit significant synteny and a conserved gene order, with approximately 1023.74 Mb (about 80.92%) of the genomic regions showing synteny between the two haplotypes(Fig.2A and Supplementary Fig. 9). However, structural and local genomic variations exist between the two haplotype genomes, encompassing 52.64 Mb of rearranged regions and 122.21 Mb of duplicated regions (Fig. 2D and Supplementary Fig. 9). These regions contain 5,091,834 SNPs, 576,235 insertions, and 445,379 deletions (Fig. 2C and Supplementary Fig. 10). Among the 101 inversion events observed, the inversion lengths range from several hundred base pairs to several mega bases, with two large inversions between Chr 1 and Chr4 exceeding 5 Mb in length. In addition, there is a 67.65 Mb unaligned region between the two haplotypes, with a segment on Chr 2 exceeding 9 Mb unique to Hap2 (Fig. 2D). This region is part of the chromosome B compartment and consists of two distinct TADs (Supplementary Fig. 11). Hi-C chromatin contact data provides strong evidence for major structural variations (SVs), further validating high accuracy of the genome assemblies and the precise identification of SVs (Fig. 2B).

**Fig. 2.**
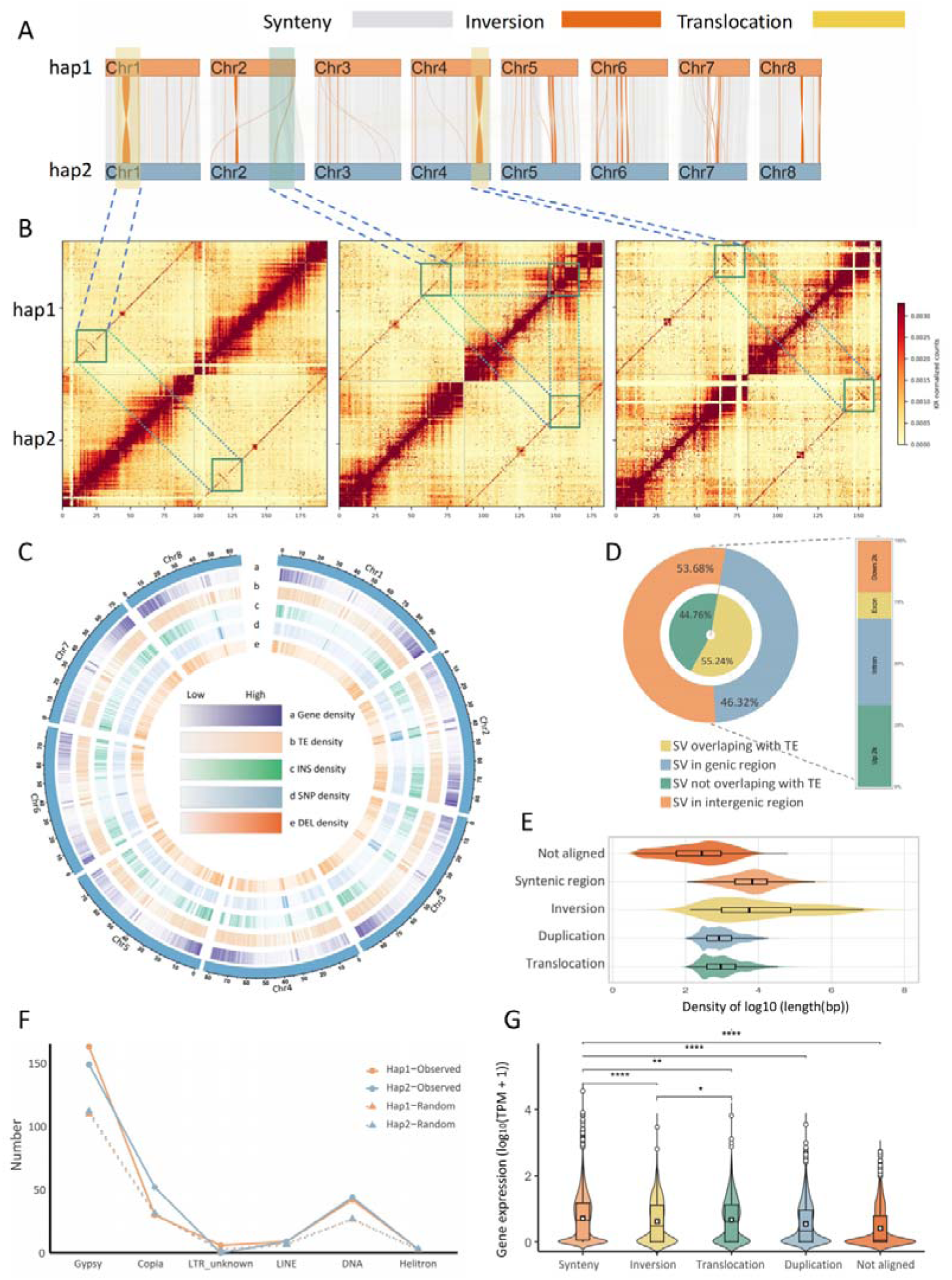
Comparison of structural variations between *I. pseudotinctoria* genome Hap1 and Hap 2, and their effects on gene expression. (A) Comparison between two haplotype genomes, with Hap1 as the reference. (B) Large inversion and deletion validation between two haplotypes. The orange regions on the collinear blocks indicate inversions. Hi-C heatmaps support these inversions and deletions. (C) Circos plots of variant distribution and frequency density, illustrating the distribution of frequency differences among haplotypes: (a) Hap1 gene density; (b) Hap1 TE density; (c) INS density; (d) SNP density; (e) DEL density. The density plots depict the distribution of variant frequencies in the reference genotype using Hap1 as the reference. (D) The proportion of SVs overlapping with TEs or different genomic regions. In the gene body, upstream 2-kb and downstream 2-kb regions of the gene body are considered genic regions and other regions are considered intergenic regions. (E) Length distribution of inter-genomic variations. (F) Counting the number of inverted breakpoints overlapping TEs in two haplotype genomes (200 base pairs per breakpoint). Solid lines represent actual counts, dashed lines represent randomized counts. (G) Violin plots showing gene expression differences across distinct variant regions. Y-axis indicates the gene expression levels of genes that overlap with the structural variants. Mann-Whitney-Wilcoxon test. *P < 0.05; **P < 0.01; ***P < 0.001; NS, no significant difference. Error bar type is the standard error (SE). The width of each violin represents the density of the data. In boxplots, the center line in the box indicates the median value, and the box height indicates the 25th to 75th percentiles of the total data. Whiskers indicate the 1.5× interquartile range. Points outside the whis kers indicate outliers.

SVs overlap with transposable elements by up to 53.68% (Fig. 2E), indicating that transposable elements (TEs) may play a significant role in the formation of structural variations. Inversions typically have a more direct and significant impact on gene expression and function. Therefore, we investigated the variation patterns of TEs with increasing distance from inversion breakpoints. The TE density distribution showed a similar overall trend between the two haplotypes relative to the inversion breakpoints. In Hap1, TE density was significantly higher near inversion breakpoints and gradually decreased with distance (R² = 0.44, p = 0.041), indicating a strong enrichment of TEs near SV sites. A similar pattern was observed in Hap2 (R² = 0.31, p = 0.074)(Supplementary Fig. 12), though the p-value was marginally significant. These results suggest that TEs are preferentially enriched near inversion boundaries and may contribute to the formation or stabilization of inversions in the *I. pseudotinctoria* genome. We further performed a randomization test by comparing the observed number of different TE types near inversion breakpoints with the number at randomly selected inversion sites. The test revealed significant enrichment of Gypsy and DNA type TEs in both haplotypes, with observed numbers significantly higher than expected from random distributions(Fig. 2F), indicating that these TE types are closely associated with structural variations at inversion breakpoints.

SVs may influence gene expression and confer haplotype-specific advantages. Among all SVs, 46.32% are located within genic and regulatory regions (±2 kb), with the majority concentrated in intronic and upstream 2kb regions, accounting for 35.09% and 33.08%(Fig. 2E), respectively. We observe that gene expression in SV regions is significantly lower than in highly collinear regions. Different SV types, such as inversions and translocations, are generally associated with reduced gene expression(Fig. 2G), while duplications correlate with higher expression levels, suggesting that gene copy number effects may substantially influence gene expression. Further comparison of gene expression in SV regions between Hap1 and Hap2 revealed that Hap1 exhibits higher gene expression in inversion and translocation regions than Hap2 across all three SV types (inversion, translocation, and duplication)(Supplementary Fig. 13). This indicates that the two haplotypes respond differently to SVs, and the effects of SVs vary across genomic regions.

### Identifying haplotype-specific and ASE genes

The characteristics of insect-pollinated flowers contribute to extensive genetic variation and allelic diversity between the two haplotypes in *I. pseudotinctoria*. To gain a deeper understanding of the origins and functional significance of these variations, we performed a detailed allelic identification between the haplotypes, following methods previously applied in pear and tea. We categorized the haplotype genes into three groups: 1) Biallelic genes (alleles with no shared coding sequence, CDS), 2) Same CDS genes (alleles with identical CDS), and 3) Haplotype-specific genes (genes present exclusively within a single haplotype) (Fig. 3A). Using this classification, we identified a total of 19,718 pairs of Biallelic genes, 5,587 pairs of same CDS genes, and 2,607 Hap1-specific genes and 2,331 Hap2-specific genes (Fig. 3B).

**Fig. 3.**
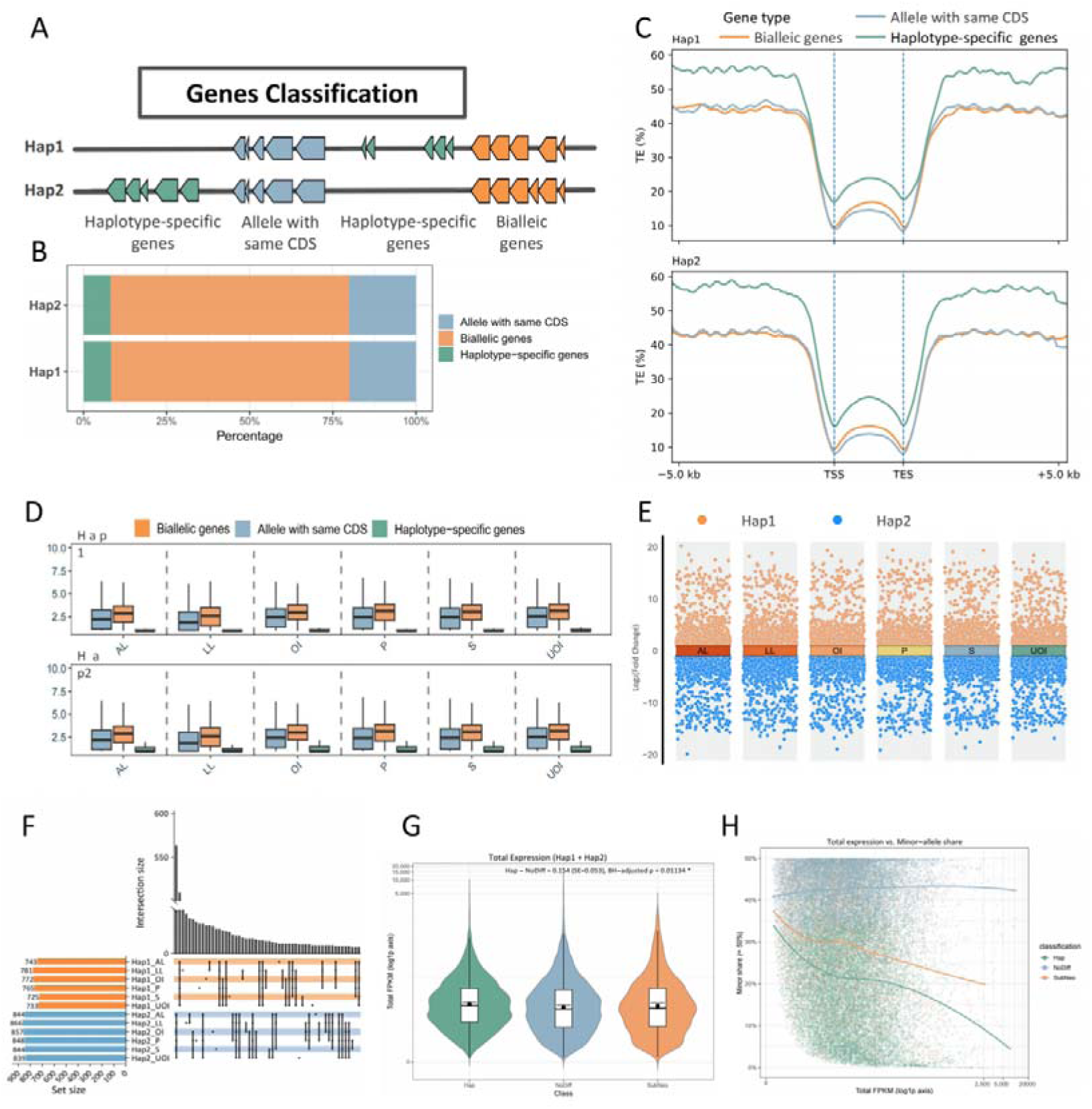
Allelic gene variation and expression pattern in *I. pseudotinctoria*. (A) The layout for allele gene classification. The alleles were divided into the haplotype-specific genes (no paired genes in the corresponding haplotype), the alleles with the same CDS and biallelic genes. (B) Bialelic proportion: The proportion of alleles within each haplotype that possess the same CDS and haplotype-specific genes. (C) Distribution of transposable elements (TEs) in the genome, as well as in the 5 kb upstream and downstream regions of the bialelic region, for alleles in Hap1 and Hap2 that share the same CDS and haplotype-specific genes. (D) The mean FPKM value of haplotype-specific genes, alleles with same CDS and bi allelic genes in Hap1 and Hap2. Apical Lateral (AL); Lateral Lateral (LL); Open Inflorescence (OI); Unopened Inflorescence (UOI); Stem (S); Pod (P). (E) The expression profile of the ASE genes in Hap1 and Hap2, P values are calculated using a two-sided quasi-likelihood F-test based on a negative binomial distribution. (F) The upset plot of ASE patterns. The horizontal bar chart on the left indicates the number of highly expressed biallelic genes within each haplotype. The vertical bar chart at the top displays the number of ASE genes within each haplotype in the sample, represented by solid dots and connected by lines. (G) Comparison of total allelic expression among ASE gene-pair classes. Violin plots show the distribution of total expression (hap1 + hap2, log1p-transformed FPKM) across ASE categories. The black dot and error bar represent the estimated marginal mean and its 95% confidence interval derived from a mixed-effect model. (H) Relationship between total allelic expression and minor-allele share across haplotype classes. Each point represents one gene pair within a specific tissue, with the x-axis indicating total FPKM (log1p-transformed) and the y-axis showing the proportion of the minor allele’s contribution to total expression. LOESS regression lines depict the smoothed trend for each ASE class (Hap, SubNeo, NoDiff).

Comparison of these gene types revealed significant differences in genomic distribution. Haplotype-specific genes showed a higher abundance of transposable elements (TEs) within their gene body and the flanking ±5kb regions, suggesting enhanced TE activity and gene regulatory potential in these regions (Fig. 3C). Furthermore, the average expression level of haplotype-specific genes across tissues was lower than that of Biallelic and same CDS genes (Fig. 3D), indicating potential differences in their regulatory mechanisms.

DNA methylation analysis in leaves revealed significantly higher methylation levels in the gene bodies and flanking (±2 kb) regions of haplotype-specific genes compared to the other gene types(Supplementary Fig. 14), suggesting stronger methylation-mediated silencing contributes to their lower expression. The presence of TEs in these regions likely plays a crucial role in this epigenetic regulation. KEGG and GO enrichment analyses revealed that Hap1 haplotype-specific genes are primarily enriched in defense- and stress-related pathways (e.g., MAPK signaling, Toll-like receptor signaling and DNA repair). In contrast, Hap2-specific genes were enriched in in pathways related to terpenoid/gingerol biosynthesis, photosynthesis, and hormone signaling (Supplementary Fig. 15). This indicates complementary functional specializations, "defense and repair" in Hap1 versus "metabolism and growth" in Hap2, that collectively strengthen the adaptability of *I. pseudotinctoria*.

We further examined the allele-specific expression (ASE) profiles across six tissues: apical lateral (AL), lateral lateral (LL), open inflorescence (OI), unopened inflorescence (UOI), stem (S), and pod (P). We identified 2,320 allelic loci with tissue-specific expression. Among these, 571 gene pairs showed consistently higher expression from Hap1, while 362 pairs displayed dominant expression from Hap2 (Fig. 3E, F). According to allelic expression patterns, ASE gene pairs were classified into three categories: haplotype-dominant (HapDom, n=1,252), sub or neo-functionalized (SubNeo, n=307), and non-differential (NoDiff, n=760). Mixed-effect modeling of total expression (Hap1 + Hap2) revealed that HapDom pairs had significantly higher overall expression than NoDiff pairs (Δ = +0.1536, SE = 0.053, p = 0.0113), indicating a dosage amplification effect. In contrast, the difference between HapDom and SubNeo pairs was not significant (p > 0.05). Analysis of minor-allele share showed significant differences in allelic bias among the three categories. HapDom pairs had a markedly lower minor-allele contribution (average 18.0%; logit difference = −1.211, p < 0.0001), compared to NoDiff pairs (42.4%). LOESS regression indicated a negative correlation between total expression and minor-allele share in HapDom pairs, suggesting that increased transcript abundance is primarily due to dominant allele amplificatio.

NoDiff pairs maintained a stable allelic ratio across expression levels, reflecting balanced bi-allelic expression (Fig. 3F and Supplementary Fig. 16). These findings indicate that the elevated total expression of HapDom pairs is driven by the combined dominant-allele amplification and minor-allele suppression, characterizing a haplotype-dominant regulatory pattern.

### Comparative genomic analysis

To investigate the phylogenetic position of *I. pseudotinctoria* and the evolution of its gene families relative to other leguminous and model plant species, we performed gene family clustering based on protein-coding genes from *I. pseudotinctoria* (Hap1) and 15 other plant species. A total of 530 gene families were identified as strict single-copy orthologs shared across all species. In *I. pseudotinctoria*, 1,326 genes were classified as unique paralogs, and 1,762 genes remained unclustered, resulting in 3,088 species-specific genes(Fig. 4 and Supplementary Fig. 18). Functional enrichment analyses of the species-specific genes were conducted. KEGG enrichment analysis suggested potential roles in hormone regulation, secondary metabolism, and stress responses, while GO enrichment analysis implied functions in environmental adaptation, root development, and defense mechanisms(Supplementary Fig. 19). To gain insight into the genome evolution of the *I. pseudotinctoria*, a phylogenetic tree was constructed based on 530 single-copy orthologous genes. The results showed that *I. pseudotinctoria* is most closely related to *C. cajan*, with an estimated divergence time of approximately 30.5 million years ago (Mya; 21.9–40.2 Mya), forming the nearest sister lineage. It clusters within the same major clade as *G. max, Vigna* spp., and *Phaseolus* spp., whereas its divergence from *L. japonicus* occurred slightly earlier (around 35 Mya). Earlier-diverging lineages such as *M. truncatula* and *P. sativum* separated approximately 50 Mya (Supplementary Fig. 20).

**Fig. 4.**
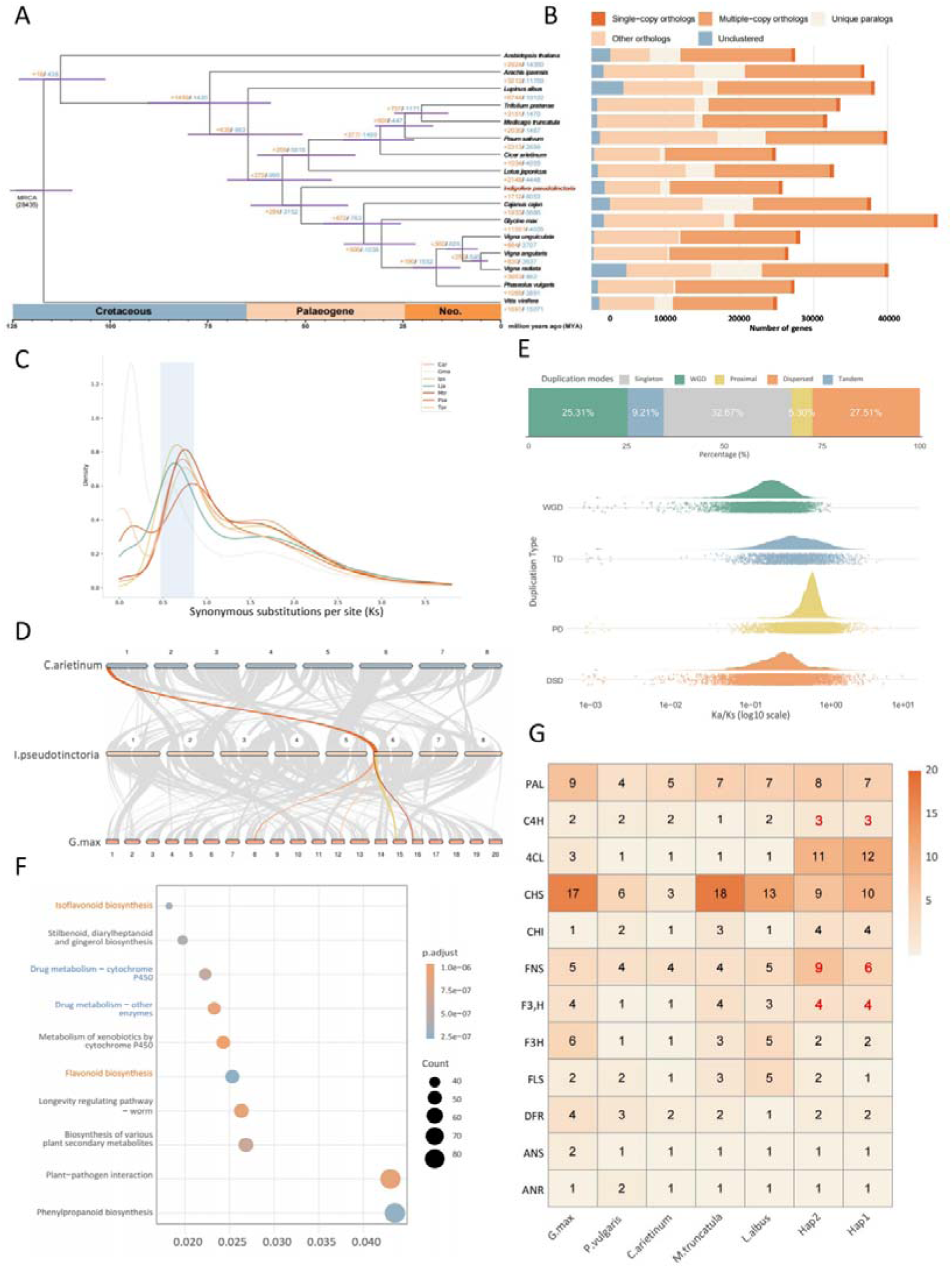
(A) Phylogenetic tree and gene family expansion/contraction based on 16 plant species. Numbers at nodes indicate divergence times. (B) Distribution of gene family numbers is shown to the right of each species. (C) Density distribution of the synonymy substitution rate (Ks) for *I. pseudotinctoria* and its closely related species. (D) Conservative gene sequence analysis of *C. cajan*, *I. pseudotinctoria*, and *G. max*. A gene fragment appears as one copy in *I. pseudotinctoria* (orange line) but four copies in G. max (red line), indicating that *I. pseudotinctoria* did not undergo a recent WGD event after legume divergence. (E) Types and proportions of duplicated genes across different models (top); cloud diagram illustrating selection pressure (Ka/Ks) on genes derived from distinct duplication patterns (bottom). (F) KEGG pathway enrichment of expanded gene families in *I. pseudotincto*ria. (G) Comparative analysis of gene copy number related to the classic flavonoid biosynthesis pathway in 6 species. Asterisks indicate gene families identified as expanded by CAFE.

Whole-genome duplication (WGD), referring to the duplication of all sequences within a genome, is a common phenomenon in plant evolution. It provides abundant genetic material for evolutionary innovation, thereby enhancing species diversity and environmental adaptability. By comparing the distribution of median synonymous substitutions (Ks) within syntenic blocks across six related species, we investigated WGD events in the *I. pseudotinctoria* genome. The main Ks peak of *I. pseudotinctoria* largely overlapped with those of *M. truncatula*, *C. cajan* and others, indicating a shared ancient legume WGD event (Ks ≈ 0.73) and the absence of any recent duplication events. In contrast, *G. max* exhibited an additional, lineage-specific WGD (Fig. 4B). Synteny analysis of the *I. pseudotinctoria* genome further supports that this species has not undergone a recent WGD event (Fig. 4C), suggesting it has retained a relatively simpler genome structure.

Gene duplication events in *I. pseudotinctoria* were categorized based on the chromosomal distribution of duplicated gene pairs, including 7064 whole-genome or segmental duplicates (WGD, 25.3%), 2571 tandem duplicates (TD, 9.2%), 1480 proximal duplicates (PD, 5.3%), and 7678 dispersed duplicates (DSD, 27.6%), with the remaining 9119 being single-copy genes (Singletons, 32.6%) (Fig. 4D). The relatively high proportion of WGD-derived genes indicates that large-scale duplication events have made substantial contributions to the expansion of the *I. pseudotinctoria* genome. To assess the evolutionary dynamics of duplicated genes, we compared the Ka/Ks ratios of gene pairs from different duplication modes. Genes derived from PDs and TDs exhibited higher Ka/Ks values than those from WGDs and DSDs (Fig. 4E), suggesting that PDs and TDs have undergone more rapid sequence divergence and potentially stronger positive selection pressure during evolution.

Gene family expansion and contraction were analyzed using Computational Analysis of Gene Family Evolution (CAFE), identifying 1,712 expanded and 8,053 contracted gene families in *I. pseudotinctoria* (Fig. 4A). GO enrichment analysis (Supplementary Fig. 21) showed that the expanded gene families were mainly enriched in functions related to responses to toxic substances, oxidoreductase activity, and stress responses. Contracted gene families were primarily involved in transmembrane transport and sesquiterpenoid biosynthetic processes. KEGG pathway analysis (Supplementary Fig. 21) further revealed that expanded families participated in pathways associated with plant–pathogen interactions and the biosynthesis of secondary metabolites, whereas contracted families were significantly enriched in oxidative phosphorylation and sphingolipid signaling pathways.

Notably, the expanded gene families were predominantly enriched in the Isoflavonoid and flavonoid biosynthesis pathway(Fig. 4F). This is particularly significant for *I. pseudotinctoria*, a medicinal plant renowned for its high flavonoid content. Consistent with this, the copy numbers of several key gene families involved in flavonoid biosynthesis, including C4H, FNS, and F3’H, were significantly increased (Fig. 4G). Furthermore, we investigated the duplication patterns of these amplified gene families within the flavonoid biosynthesis pathway, identifying tandem duplication events within the FNS and F3’H gene families. These tandem duplication events in key flavonoid biosynthetic genes are likely closely associated with the high accumulation of flavonoids characteristic in *I. pseudotinctoria*.

### Identification of flavonoid biosynthetic enzyme genes

To systematically characterize the distribution patterns of secondary metabolites across various tissues of *I. pseudotinctoria*, we employed a widely targeted metabolomics approach to profile and quantify metabolites in six distinct tissues: apical lateral (AL), lateral lateral (LL), open inflorescence (OI), unopened inflorescence (UOI), stem (S), and pod (P). A total of 2,766 metabolites were detected and classified into 13 major categories (Fig. 5B). Among these, flavonoids represented the predominant class, accounting for 25.71% of the total metabolites, followed by amino acids and their derivatives (10.39%), terpenoids (10.06%), and phenolic acids (8.41%). The dominance of flavonoids within the metabolic network of *I. pseudotinctoria* highlights the species’ remarkable metabolic specialization in flavonoid biosynthesis and accumulation. After normalization, hierarchical clustering of metabolomic data revealed distinct metabolic accumulation patterns among tissues (Fig. 5A). Overall, flavonoid compounds were highly abundant in multiple tissues, with particularly high levels in pods and floral organs (OI and UOI), whereas their levels were relatively lower in the stem, apical lateral (AL) and lateral lateral (LL). To further elucidate the biosynthetic mechanisms underlying flavonoid accumulation, RNA-seq analysis was conducted on the same six tissues of *I. pseudotinctoria*. In total, 133.89 GB of raw reads were generated, with 95.17–96.74% of clean reads successfully mapped to the Hap1. The correlation coefficients among three biological replicates exceeded 0.98 (Supplementary Fig. 22), and principal component analysis (PCA) showed tight clustering of replicates (Supplementary Fig. 23), supporting the high reproducibility of the transcriptomic data.

**Fig. 5.**
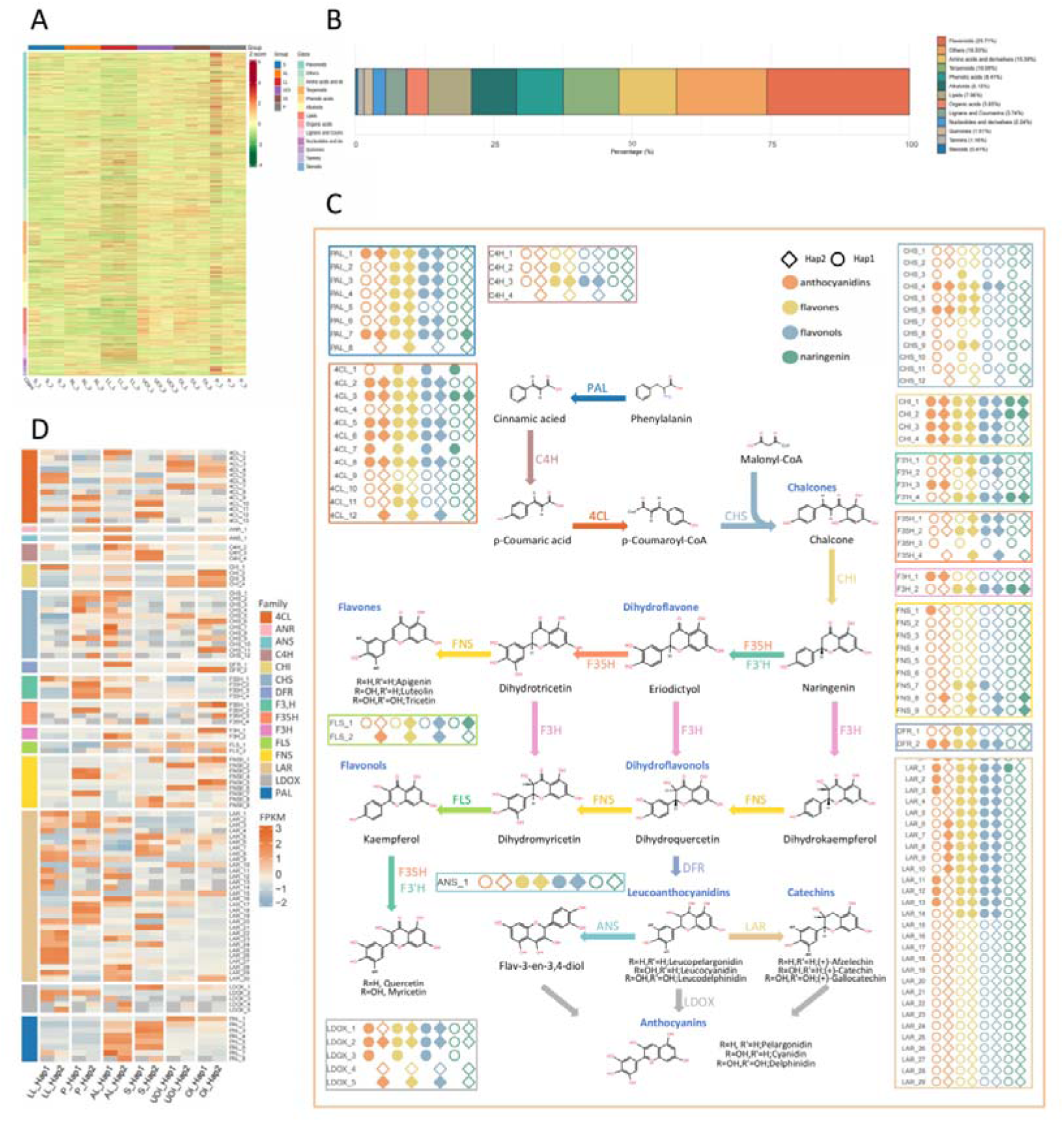
Expression analysis of enzyme-coding genes involved in flavonoid biosynthesis in *I. pseudotinctoria*. (A) Standardized abundance (Z-score) of metabolites detected in various tissues. (B) Proportion of metabolite classes detected. (C) Heatmap of expression levels of genes encoding enzymes involved in flavonoid synthesis across various tissues. (D) Biochemical pathway diagram of *I. pseudotinctoria* flavonoids, coloured graphics represent specific metabolites.

Based on our gap-free haplotype-resolved genome and previously characterized enzymatic genes involved in flavonoid pathways, we reconstructed the putative flavonoid biosynthetic network in *I. pseudotinctoria* (Figs. 5C and 5D). The pathway comprises upstream enzymes (PAL, C4H, 4CL), core structural genes (CHS, CHI, F3H, F3’H), downstream reductases and modification enzymes (DFR, ANS, LAR, LDOX), and several branch-specific enzymes (FNS, FLS). Consistent with metabolomic results, most structural genes displayed markedly higher expression in pods. Upstream, midstream, and downstream genes showed strong expression signals in pods (P), indicating that flavonoid biosynthesis is most active in this tissue. A subset of these genes also displayed moderate to high expression in open inflorescence (OI) and unopened inflorescence (UOI), though at slightly reduced levels, suggesting that floral tissues maintain elevated flavonoid fluxes but to a lesser extent than pods. In contrast, most genes were weakly expressed in leaves (LL and AL) and stems (S), with only a few upstream families (PAL, C4H, 4CL) showing relatively higher expression. Both haplotypes (Hap1 and Hap2) displayed similar overall expression patterns, indicating that alleles from both haplotypes jointly regulate flavonoid metabolism in *I. pseudotinctoria*.

To refine candidate gene identification, we integrated metabolomic and transcriptomic data to assess correlations between gene expression levels and four major classes of secondary metabolites—anthocyanidins, flavones, flavonols, and naringenin (Fig. 5D). Spearman correlation coefficients were computed for all gene–metabolite pairs, and multiple testing corrections were performed using the Benjamini–Hochberg method (FDR < 0.05, |r| ≥ 0.8). Most of key structural genes, particularly those from the PAL, 4CL, CHS, CHI, LAR, and LDOX families, showed significant correlations with at least one flavonoid subclass, supporting their cooperative roles in the branched flavonoid metabolic network of I. pseudotinctoria. Upstream genes (PAL, 4CL) exhibited broad-spectrum correlations with both flavones and flavonols, whereas midstream genes (CHS, CHI, F3H, F3’H) demonstrated branch-specific enhancements. Downstream enzymes (DFR, LDOX, LAR) were strongly correlated with anthocyanidins, implying their key roles in terminal pigment formation. Within the PAL family, multiple members in both haplotypes were significantly correlated with flavones and flavonols, with PAL_7 also showing significant correlations with naringenin and anthocyanidins, reflecting its function as a flux-control point in the pathway. Members of the 4CL family (notably 4CL_2, 4CL_3, 4CL_5, 4CL_6, 4CL_8) exhibited strong correlations with nearly all metabolite classes, suggesting broad substrate specificity and role in precursor supply for multiple branches. In the CHS family, CHS_4 and CHS_6 in both haplotypes were significantly correlated with anthocyanidins, flavones, and flavonols, implying their major contribution to flavonoid scaffold formation. All four CHI genes (CHI_1–CHI_4) were significantly correlated with all four metabolite categories, supporting their central catalytic role in the midstream reactions. The F3H and F3’H families showed clear branch preferences: F3H_1 and F3H_2 correlated strongly with flavonols and naringenin in both haplotypes, whereas F3’H genes (especially F3’H_1 and F3’H_3) were mainly correlated with flavones and flavonols, suggesting roles in hydroxylation diversity and product spectrum determination. Finally, DFR_2 and LDOX_1–LDOX_3 were significantly correlated with anthocyanidins, flavones, and flavonols, identifying them as key catalysts in anthocyanidin biosynthesis.

### Detection of flavonoid biosynthetic TFs in I. pseudotinctoria

Building upon previous phytochemical investigations that identified diverse flavonoid compounds from I. pseudotinctoria, this study aimed to elucidate the transcriptional regulatory mechanisms underlying their biosynthesis. To achieve this, we integrated metabolomic profiling with transcriptomic network analyses to identify key metabolites, co-expression modules, and regulatory genes associated with flavonoid accumulation. Firstly, five representative flavonoid compounds including Calycosin, Butein, Sulfuretin, Chrysoeriol, and Genistin, were selected for further analysis (Fig. 6A; Supplementary Table 8) based on their high abundance and strong variable importance in projection (VIP) scores (VIP > 1, P < 0.05) in the current metabolomic dataset, as well as their consistent detection in previous phytochemical studies. Together, they represent the major structural subclasses of isoflavones, chalcones, and flavonoid glycosides, serving as key biochemical indicators for subsequent transcriptional regulatory analyses.

**Fig. 6.**
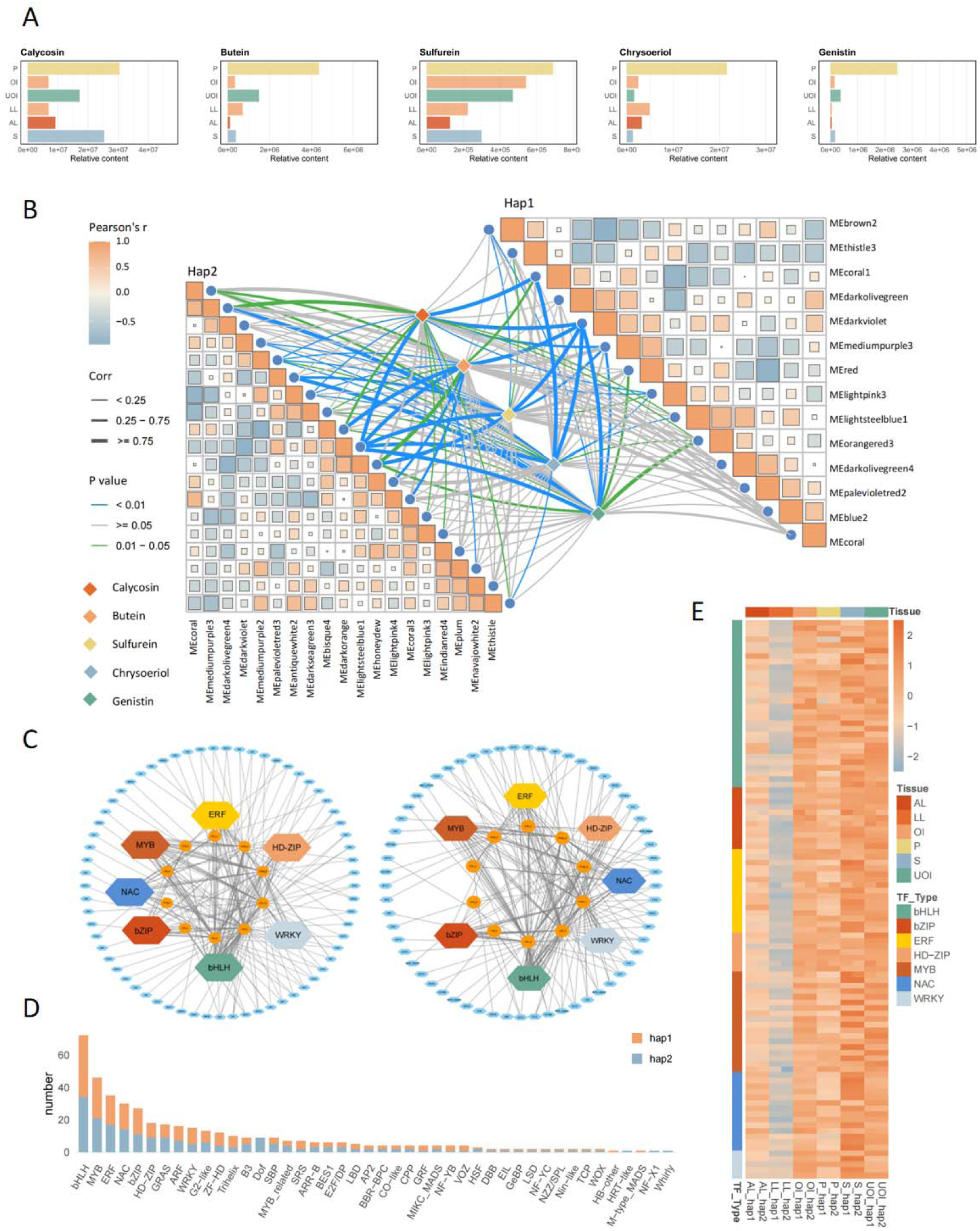
Comparative analysis of isoflavonoid contents and their regulatory network in different tissues. (A) Relative contents of five major isoflavonoids (calycosin, butein, sulfurein, chrysoeriol, and genistin) across tissues. (B) Correlation analysis and module–metabolite associations. (C) Transcription factor co-expression networks for Hap1 and Hap2 modules. (D) Distribution of transcription factor families between Hap1 and Hap2. (E) Heatmap showing transcription factor expression patterns across tissues.

Secondly, to elucidate the regulatory mechanisms underlying the biosynthesis of the five selected flavonoids, we performed a weighted gene co-expression network analysis (WGCNA), which retained 19,536 (Hap1) and 19,344 (Hap2) highly expressed genes that were further grouped into 14 and 19 co-expression modules, respectively, based on gene expression correlations and hierarchical clustering (Supplementary Fig. 24). Several modules showed strong correlations with the five flavonoid metabolites. Notably, the MEdarkviolet (Hap1) and MEdarkorange (Hap2) modules exhibited significant positive correlations with Calycosin, Butein, Sulfuretin, Chrysoeriol, and Genistin (r > 0.75, p < 0.01), indicating that genes in these modules may play crucial roles in the biosynthesis of these flavonoids. Further enrichment analysis revealed that 27 (Hap1) and 32 (Hap2) genes within these two modules were significantly enriched in KEGG pathways related to flavonoid biosynthesis. These candidate genes were mapped onto a reconstructed flavonoid biosynthesis pathway model (Fig. 5C). To eliminate potential false positives arising from haplotype-specific expression, we performed allele co-occurrence screening between the two haplotypes and retained only biallelic genes present in both Hap1 and Hap2. This filtering identified 10 high-confidence candidate genes common to both haplotypes were identified (Supplementary Table 9). These genes displayed distinct tissue-specific expression patterns across different tissues and haplotypes (Supplementary Fig. 25). Specifically, CHS, FNS, and LAR were highly expressed in open inflorescences (OI) and stems (S), whereas PAL and 4CL showed moderate and broad expression across all tissues, consistent with their catalytic roles in the early stages of phenylpropanoid metabolism. In contrast, most candidate genes exhibited relatively low expression levels in pods (P), suggesting a temporal delay between metabolite accumulation and feedback regulation that may in a metabolic equilibrium characterized by high accumulation but low transcription.

To further dissect the transcriptional regulation underlying flavonoid biosynthesis, we constructed a comprehensive candidate gene–transcription factor (TF) co-expression network based on the WGCNA modules that showed significant correlations with the five representative flavonoids (r > 0.85, P < 1E–10) (Fig. 6C). These TFs primarily belong to the MYB, bHLH, NAC, WRKY, HD-Zip, ERF, and bZIP families (Fig. 6D; Supplementary Table 10), all of which are known to participate in the regulation of secondary metabolism. To support the co-regulatory hypothesis, we predicted cis-acting elements in the promoter regions of the candidate genes and identified binding sites for these TF families (Supplementary Fig. 26). These findings provide compelling evidence for hierarchical transcriptional regulation between TFs and enzyme-encoding genes involved in flavonoid biosynthesis. Notably, these TFs exhibited distinct tissue-specific expression patterns with the higher expression levels in stem (S) and open inflorescence (OI) but lower levels in leaves (Fig. 6E). Interestingly, this spatial expression pattern is not concordant with the distribution of flavonoid metabolites, which predominantly accumulate in pods. This discrepancy implies a spatial decoupling between transcriptional regulation and metabolite deposition, suggesting that flavonoid biosynthesis is transcriptionally activated in early or structural tissues (e.g., S and OI), while the pods mainly function as storage organs for accumulation of these compounds, potentially through inter-tissue transport or feedback-controlled redistribution.

### Expression of flavonoid genes during colour change in flowers

*I. pseudotinctoria* possesses not only significant medicinal value but also considerable ornamental potential, largely due to the vivid colouration of its flowers, a trait closely associated with anthocyanin biosynthesis. Metabolomic analysis identified 29 anthocyanins and their derivatives, with several compounds showing markedly higher accumulation in open inflorescences (OI) compared with other tissues, particularly those directly involved in colour formation (Fig. 7A). Principal component analysis (PCA) further revealed clear tissue separation, indicating distinct spatial patterns of anthocyanin metabolism (Fig. 7B). Among these metabolites, delphinidin-3-O-rutinoside-7-O-glucoside and delphinidin 3,5,3′-triglucoside were enriched in UOI, suggesting the predominance of bluish-purple anthocyanins at early floral stages. In contrast, petunidin-3,5-di-O-glucoside, cyanidin-3-O-glucoside, malvidin-3-O-arabinoside, malvidin-3-O-glucoside, delphinidin 3,5-diglucoside, and malvin accumulated predominantly in OI tissues, implying enhanced methylation activity during flower opening and a progressive colour transition from bluish-purple to purplish-red (Fig. 7B, C). To elucidate the molecular basis of these metabolic shifts, the anthocyanin branch of the flavonoid biosynthetic pathway was reconstructed in I. pseudotinctoria to identify key enzymatic genes (Fig. 7D). In this pathway, leucoanthocyanidin acts as a central intermediate that is converted by ANS and LDOX into cyanidin, which is subsequently modified by F3′5′H and OMT to yield delphinidin, petunidin, and malvidin. These pigments are then stabilized through glycosylation by UGT enzymes to form derivatives such as malvin. Multiple OMT and UGT gene candidates were identified, with OMTs showing strong expression during late flower development and UGTs acting primarily in terminal glycosylation steps. The coordinated upregulation of OMT genes alongside increased accumulation of methylated anthocyanins highlights their pivotal role in pigment diversification and the dynamic modulation of flower colouration in I. pseudotinctoria.

**Fig. 7.**
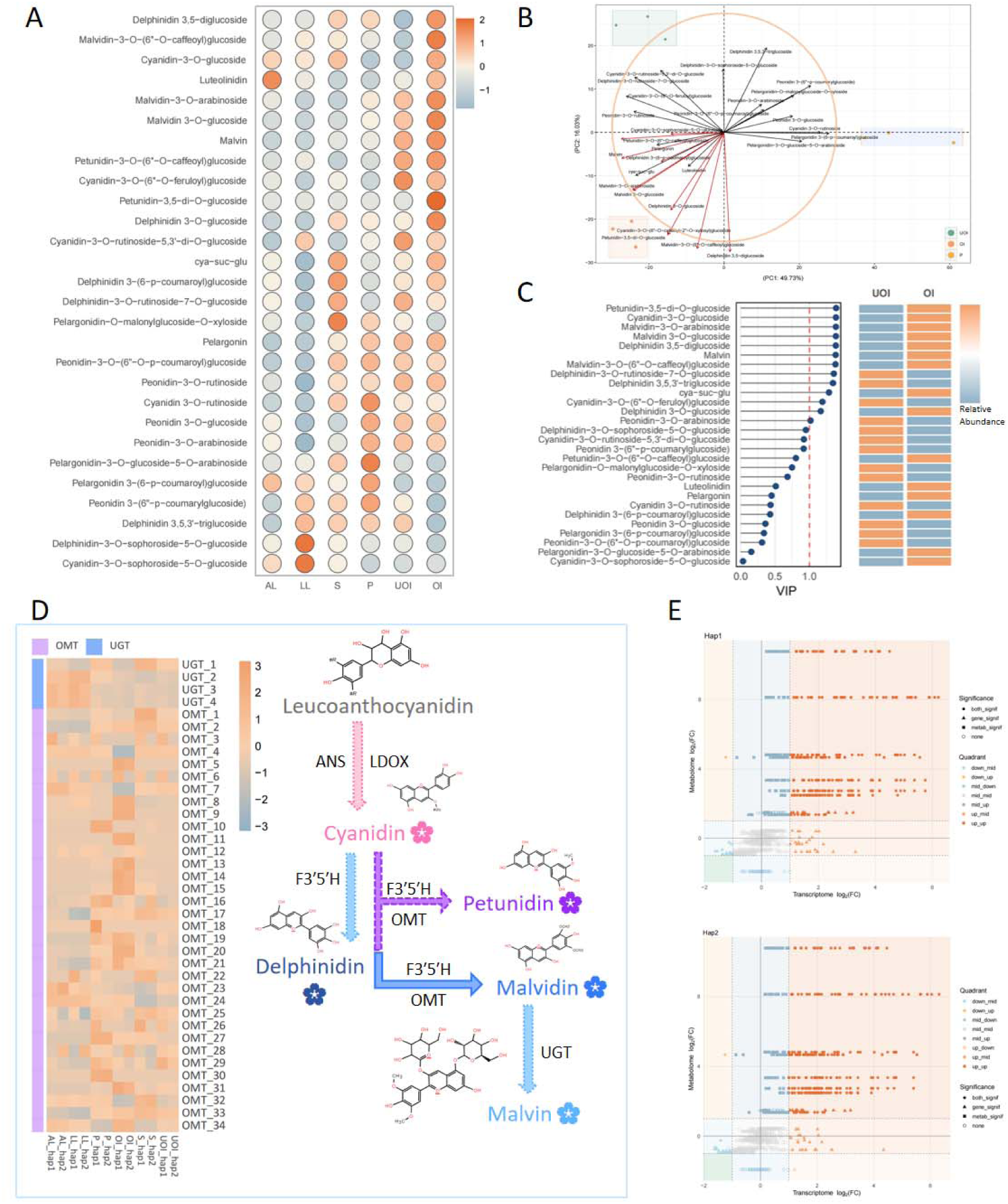
Metabolic and transcriptional profiling of anthocyanin-related compounds across different tissues. (A) Heatmap depicting the relative abundance of anthocyanin derivatives across different tissue samples (AL, LL, S, P, UQI, OI). (B) Principal component analysis (PCA) plot showing the clustering of tissue samples based on their metabolite composition. (C) VIP (Variable Importance in Projection) score analysis of anthocyanins and their tissue-specific abundance. (D) Gene expression profiling of key enzymes involved in anthocyanin biosynthesis, including OMT and UGT, with the corresponding metabolic pathways of key anthocyanins.

Integrated transcriptomic and metabolomic analyses of two Hap further revealed coordinated regulation of genes and metabolites (Fig. 7E; Supplementary Table 11). A nine-quadrant comparison of log□(fold change) values demonstrated a strong positive correlation between transcriptional activity and metabolite abundance, with most upregulated pairs clustering in the up_up quadrant. Pathway annotation and differential expression analyses identified ten key genes associated with anthocyanin biosynthesis, mainly belonging to the OMT and F3′5′H families. Among them, OMT_8, OMT_9, OMT_20, OMT_31, OMT_34, and F3′5′H_1 were upregulated in both haplotypes, while OMT_6 and OMT_11 were Hap1-specific, and OMT_13 and F3′5′H_3 were elevated only in Hap2.

## Discussion

In this study, we generated a gap-free, T2T, haplotype-resolved reference genome for *I. pseudotinctoria*, representing the first complete genome assembly within the genus Indigofera. This high-quality assembly establishes a foundational genomic resource for the genus and marks a significant milestone that will facilitate future investigations into phylogenomics, trait evolution, biosynthetic pathway elucidation, and adaptive biology across Indigofera and the wider Fabaceae. By integrating PacBio HiFi sequencing, Oxford Nanopore ultra-long reads, and Hi-C chromosomal scaffolding, we obtained two highly contiguous and accuracy-validated haplotype assemblies, each exhibiting near-complete chromosome-length continuity, high LAI values, and excellent QV scores. Moreover, the complete recovery of all telomeric and centromeric regions supported that both haplotypes extend to the authentic chromosomal termini. Together, this T2T genome provides a crucial, high-resolution genomic framework that will substantially enhance functional gene discovery and evolutionary analyses in Indigofera pseudotinctoria and its closely related lineages.

*I. pseudotinctoria* is a perennial plant with insect-pollinated and highly outcrossing hybridity. Despite extensive homology between the two haplotypes, significant structural variations such as inversions, duplications, and translocations highlight the dynamic character of its genome. The strong co-localization of these structural variants with TEs suggests that TEs act as likely contribute of large-scale genomic inversions. The enrichment of Gypsy and DNA transposons near inversion breakpoints supports their potential involvement in the formation or stabilization of these variants. Such patterns mirror TE-mediated chromosomal evolution reported in other legume species^35^, underscoring the conserved role of mobile elements in shaping plant genome diversification^36^.

Asymmetric allelic expression provides further evidence for functional divergence between the two haplotypes^37^. Nearly half (46.32%) of the structural variants overlap with genic or regulatory regions, where expression levels are significantly reduced compared with collinear loci, suggesting that local structural context may contribute to transcriptional repression. The elevated transposable element density and enhanced DNA methylation observed in haplotype-specific genes indicate that structural variation and epigenetic silencing act synergistically to reinforce regulatory asymmetry, implicating epigenetic repression in haplotype-specific regulation^29,31,38^. The functional specialization of the two haplotypes is also non-random: Hap1 is enriched for defense- and stress-related pathways, whereas Hap2 predominantly contributes to metabolic and photosynthetic processes. This division of regulatory labor may provide environment-dependent flexibility, enabling rapid sHiFits between growth and stress-adaptive states. Allele-specific expression (ASE) analyses revealed tissue-dependent dominance patterns. ASE patterns further reveal widespread haplotype dominance, where one allele is consistently upregulated while its counterpart is suppressed. Such coordinated expression bias suggests a dosage-balancing mechanism that fine-tunes transcript output and supports phenotypic plasticity. Such regulatory asymmetry may underlie dosage compensation and phenotypic plasticity in *I. pseudotinctoria*, consistent with observations in pear^29^ and apple^39^. The coexistence of pronounced structural heterozygosity, TE-driven genome plasticity, and allelic expression divergence could highlight the complex interplay between genome architecture and regulatory evolution in Indigofera pseudotinctoria.

Comparative genomic analyses indicate that Indigofera pseudotinctoria occupies a unique phylogenetic position within the legume family, forming a sister clade to *Cajanus cajan* and diverging approximately 30.5 Mya. After the ancient legume-wide duplication event (∼Ks = 0.73), this lineage did not undergo any subsequent lineage-specific whole-genome duplication (WGD), highlighting the genomic stability of the Indigofera genus and distinguishing it from *Glycine max*^40^. Instead, evolutionary innovation in *I. pseudotinctoria* appears to have arisen through localized duplications and subsequent neofunctionalization, particularly via tandem and proximal duplications that exhibit elevated Ka/Ks ratios and signatures of positive selection. Expansion of gene families associated with oxidoreductase activity, plant–pathogen interactions, and secondary metabolism, accompanied by contraction of those related to energy metabolism, reflects adaptive genomic reorganization toward enhanced defense and stress tolerance. Notably, the significant expansion of key gene families involved in flavonoid and isoflavonoid biosynthesis (including C4H, FNS, and F3’H) corresponds with the exceptionally high accumulation of bioactive flavonoids characteristic of *Indigofera* species.^8,41^. These tandem duplications may have conferred heightened metabolic plasticity, facilitating both ecological adaptation and the diversification of medicinal properties in *I. pseudotinctoria*.

The integration of metabolomic and transcriptomic analyses revealed that Indigofera pseudotinctoria exhibits a pronounced specialization toward flavonoid biosynthesis, reflecting both ecological adaptation and the biochemical foundation of its medicinal properties. Among the 2,766 metabolites detected, five representative flavonoids, including Calycosin, Butein, Sulfuretin, Chrysoeriol, and Genistin, were highly enriched in pods and serve as key bioactive markers due to their recognized pharmacological functions. Calycosin and Genistin, two major isoflavones, possess strong anti-inflammatory, estrogenic, and vasoprotective activities^42,43^, while the chalcones Butein and Sulfuretin exhibit antioxidant and anti-tumor capacities. Chrysoeriol, a methylated flavone with neuroprotective and anti-diabetic properties, further emphasizes the pharmacological breadth of this species^44^. The preferential accumulation of these compounds in pods suggests that reproductive tissues act as metabolic reservoirs for pharmacologically active flavonoids, consistent with their roles in plant defense and reproductive fitness^45^, and aligning with the long-standing use of pod extracts in traditional medicine. These findings invite an evolutionary ecological interpretation. Previous studies have shown that many plants enhance their tolerance to poor and nutrient-deficient soils, as well as to cold, UV-B radiation, and drought, by upregulating flavonoid and phenylpropanoid biosynthesis^46–49^. These compounds serve as photoprotectants, antioxidants, and membrane stabilizers under environmental extremes. Such environmentally induced metabolic enrichment not only confers physiological resilience but also likely shaped the evolutionary trajectory of the plants. The ability to synthesize and store large amounts of flavonoids may thus have provided *I. pseudotinctoria* with a selective advantage in arid and high-irradiance habitats, indirectly driving the enhancement of its medicinal properties.

Currently, research into the developmental and formation mechanisms of flavonoid synthesis in *I. pseudotinctoria* remains relatively limited. WGCNA analysis has identified several core structural genes involved in synthesising key flavonoid metabolites in the *I. pseudotinctoria*, including PAL, 4CL, CHS, CHI, F3H, F3′5′H, DFR, and LDOX, alongside multiple transcription factors (TFs) participating in the biosynthesis of these compounds. MYB, bHLH, NAC, WRKY, and ERF TF families were found to be strongly co-expressed with these structural genes, suggesting a hierarchically coordinated transcriptional network. This inference is bolstered by functional studies in model species, where MYB TFs such as AtMYB12 in Arabidopsis^50^ and GmMYB29 in *Glycine max*^51^ activate flavonoid biosynthesis genes, including CHS, CHI, and F3H, while bHLH TFs like TT8 and GL3 form MYB-bHLH-WD40 (MBW) complexes that regulate tissue-specific pigmentation and stress-induced flavonoid accumulation^52^. Furthermore, one striking observation in our results is the spatial mismatch between gene expression and metabolite accumulation: key enzyme genes and regulatory transcription factors displayed the highest expression levels in stems and open inflorescences, whereas the five downstream metabolites accumulated most abundantly in pods. This pattern indicates a spatial decoupling between flavonoid biosynthesis and accumulation, revealing a source–sink metabolic strategy rather than a strictly tissue-autonomous biosynthetic mechanism. This phenomenon may reflect a developmentally coordinated metabolic partitioning mechanism in *I. pseudotinctoria*. Stems and inflorescences, as metabolically active tissues, possess high precursor fluxes from primary metabolism, enabling de novo synthesis of flavonoid aglycones driven by the transcriptional activation of structural genes and transcription factor networks. In contrast, pods may function as long-term storage organs, accumulating glycosylated or modified flavonoids that are transported through phloem loading or cellular transporters^53^. Moreover, the “high accumulation low transcription” equilibrium observed in pods suggests a feedback regulation mechanism, flavonoid deposition may locally suppress biosynthesis to avoid metabolic overload, sHiFiting dependence on precursor supply from upstream tissues^54^. This model aligns with recent findings in Medicago truncatula^55^, Glycine max^56^, and *Camellia sinensis*^57^, where specialized metabolites are synthesized in vascular or reproductive precursor tissues and subsequently transported to storage organs. In *I. pseudotinctoria*, such a long-distance allocation mechanism may partially explain its medicinal efficacy, pharmacologically active flavonoids are actively mobilized into pods, which corroborates the traditional preference for using pod extracts in herbal medicine^58,59^.

## Materials and methods

### Plant materials and DNA extraction

Genomic DNA was isolated from a single plant of *Indigofera pseudotinctoria* cv.’Yudong’ (Germplasm ID: XKY20150201), which was developed through multiple cycles of recurrent mass selection using progenitor germplasm collected from Wushan County, Chongqing City, China. Young leaf samples were flash-frozen in liquid nitrogen, and high-molecular-weight DNA was extracted using the SDS method followed by purification with a QIAGEN® Genomic-tip kit (Cat. #13343). DNA integrity was verified by electrophoresis on 0.75% agarose gels, with degradation monitored by fragment size distribution. Purity was assessed using a NanoDrop™ One spectrophotometer (Thermo Fisher Scientific), confirming absorbance ratios of A260/280 = 1.8–2.0 and A260/230 = 2.0–2.2. Final quantification was performed with a Qubit® 3.0 Fluorometer (Invitrogen) and the dsDNA HS Assay Kit.

### Library preparation and sequencing

PacBio HiFi SMRTbell libraries were constructed following the standard protocol using the SMRTbell Express Template Prep Kit 2.0 (PacBio, CA, USA). The lengthy DNA fragments were skillfully sheared down to 15-18 kb using a g-TUBE (Covaris, MA, USA). Single-strand overhangs were cut off, and damaged and broken DNA was patched up with the chemicals in the Template Prep Kit. Once the ends were fixed, SMRTbell hairpin adapters were ligated to them, and then the libraries were concentrated and purified using AMPure PB beads (PacBio, CA, USA). BluePippin was utilized to size-select SMRTbell templates more than 15 kb to get large-insert SMRTbell libraries for sequencing (SageScience, MA, USA). The sequencing was performed using a PacBio Sequel II device with Sequencing Primer V2 and a Sequel II Binding Kit 2.0. For the raw sequencing reads, the min passes = 3 and min RQ = 0.99 default parameters in CCS software (https://github.com/PacificBiosciences/ccs) were utilized to generate high-precision HiFi reads with quality over Q20.

For ONT sequencing, the NEBNext Ultra II End Repair/dA-tailing Kit was used to fix the ends of the lengthy DNA fragments that were size-selected with the BluePippin system (Sage Science, USA). The fragment size of the library was then measured with a Qubit® 3.0 Fluorometer after a second ligation reaction was performed with an LSK109 kit (Invitrogen, USA). Library sequencing was performed on Nanopore PromethION instruments (ONT: Oxford Nanopore Technologies, UK). The raw information is presented as FAST5 binary signal data. We utilized a high-precision flip-flop model with the guppy basecaller command in the GPU-enabled Guppy program (v3.4.4) to collect the fastq data. Reads with Q scores greater than 7 were considered passed after the raw data in fastq format had been analyzed for base quality.

In addition, a Hi-C library was constructed and sequenced to facilitate chromosome level genome assembly. Approximately 2 g of fresh leaves were utilized for library construction, and the technique involved formalin fixation, crosslinking, nuclei suspension, digestion, DNA ligation, end-repair, purification, and quantification, as previously described. The qualifying library was subsequently sequenced on an MGI-2000 platform. The quality control measures were identical to those described above for paired-end sequencing

### Genome assembly

High-molecular-weight genomic DNA of Indigofera pseudotinctoria was sequenced using a combination of PacBio HiFi, Oxford Nanopore Technologies (ONT) ultra-long, and Hi-C sequencing technologies to achieve a telomere-to-telomere (T2T), haplotype-resolved assembly.

#### (1) De novo haplotype-resolved assembly

Raw PacBio HiFi reads were used as the backbone for initial de novo assembly, with ONT ultra-long reads providing long-range continuity across repetitive regions. The HiFi, ONT and Hi-C datasets were co-assembled using HiFiasm^32^ in haplotype-resolved mode, generating two phased assemblies (Hap1 and Hap2). This process leveraged heterozygous variant information to partition reads into haplotype-specific bins before contig construction. Contigs showing non-nuclear origin (microbial or plastid contamination) were filtered out, yielding pure haplotype contigs.

#### (2) Chromosome anchoring and scaffolding

Hi-C data were then integrated to scaffold each haplotype independently using Haphic^60^, generating chromosome-scale assemblies. Hi-C contact matrices were visualized in Juicebox^61^, and scaffolds were manually corrected for misjoins or inversions to ensure accurate chromosomal phasing.

#### (3) Gap filling and telomere-to-telomere completion

To close remaining gaps, ONT ultra-long and PacBio HiFi reads were reassembled using HiCanu^33^ and NextDenovo^34^, followed by iterative gap-filling with TGS-GapCloser^62^ and quarT2T^63^. Gap regions identified on chromosomes 1, 5, and 8 were manually curated and verified through read remapping. The telomeric region was identified by screening for the characteristic plant telomeric repeat sequence AAACCCT. CentIERv2.0^64^ is used for centromere identification, and merge the centromere sequences obtained by the two methods.

### Genome annotation

#### (1) Repeat annotation and TE analysis

Repetitive elements of the *I. pseudotinctoria* genome were identified using a combination of structure-based, de novo, and homology-based approaches. Long terminal repeat (LTR) retrotransposons were detected using LTR_harvest^65^, LTR_FINDER^66^, and refined with LTR_retriever^67^. De novo prediction of other transposable elements was performed using RepeatModeler^68^. Homology-based annotation was carried out using RepeatMasker^69^ with the Dfam^70^ and RepBase^71^ databases. The resulting libraries were merged, filtered for protein-coding contamination using ProtExcluder^72^, deduplicated with VSEARCH^73^, and further classified with DeepTE^74^. Tandem repeats were identified using TRF^75^, and protein-level repeats were detected with RepeatProteinMask^76^.

#### (2) Non-coding RNA annotation

Non-coding RNAs (ncRNAs) were identified using a combination of structure- and homology-based approaches. Small RNAs including snoRNAs, snRNAs, and other Rfam families were detected using Infernal^77^ against the Rfam database with parameters --rfam --cut_ga --nohmmonly. Ribosomal RNAs were predicted using RNAmmer^78^ with the eukaryotic model. Transfer RNAs (tRNAs) were identified using tRNAscan-SE^79^ in eukaryotic mode (-E -G), generating both structural and statistical outputs.

#### (3) Genes annotation

For gene prediction in a TE-masked genome, three approaches were utilized: ab initio prediction, homology-based search, and transcriptome-guided identification. BRAKER^80^ was employed for homologous peptide alignment from closely related species to the genome assembly, providing gene structure information for homology-based prediction. RNA-seq-based gene prediction involved aligning filtered mRNA-seq reads to the reference genome using HISAT2^81^, followed by transcript assembly with StringTie^82^. For transcriptome-guided prediction, RNA-seq reads were assembled using StringTie to generate a training set, which was used with Augustus for ab initio gene prediction. Finally, TSEBRA ^80^was used to integrate the gene set. Untranslated regions (UTRs) and alternative splicing regions were defined using GUSHR^83^ and GeMoMa^84^ based on RNA-seq alignments, and the longest transcripts for each locus were retained, with regions outside the open reading frames (ORFs) designated as UTRs.

### Haplotype comparison and structural variation analysis

Synteny between Hap1 and Hap2 was assessed using MUMmer4^85^ and visualized with GenomeSyn^86^. Structural variationsm were identified by SyRI^87^. Trandomization anhe overlap between inversion regions and transposable elements was calculated to infer TE-mediated rearrangements. The distances between inversion regions and their nearest transposable elements (TEs) on each chromosome were calculated using BEDTools^88^. Here, TEs included both observed and randomly generated elements. Random TEs were simulated by shuffling the positions of inversion regions 100 times across the genome to establish a null expectation. Expression levels of genes within SV regions were compared using RNA-seq FPKM data to evaluate transcriptional impacts.

### Identification of haplotype-specific and allele-specific expression (ASE) genes

Allelic gene pairs between the two *I. pseudotinctoria* haplotypes were identified following approaches previously described in tea and pear. Two complementary strategies were applied: synteny-based and coordinate-based allele identification. Syntenic blocks were identified using JCVI^89^ with the parameter --cscore=0.99, and genes located within one-to-one syntenic regions were considered putative allelic pairs. For genes not located within syntenic blocks, homologous relationships were determined using Diomand^90^ (E-value < 1e−5, identity >80%, coverage >80%) and GMAP^91^ based on genomic coordinate overlap (>50%). Genes with identical coding sequences (CDS) were defined as same CDS genes, while unpaired genes were classified as haplotype-specific genes. Further allele-specific gene expression analysis was conducted on the Biallelic genes. Methylation levels from Oxford Nanopore sequencing reads were extracted using Nanopolish^92^, and BEDTools^88^ was used to analyze the distribution of transposable elements (TEs) and methylation levels across different gene categories.

To investigate the regulatory mechanisms of allelic gene expression, RNA sequencing data from six tissues were mapped to the haplotype-resolved genome using HISAT2^81^. FPKM estimates were calculated using StringTie^82^ based on uniquely mapped reads. Allele-specific expression (ASE) was assessed using DESeq2^93^, with a fold change greater than 2 and a corrected P-value < 0.05. Following previous studies, ASE genes were further classified into three categories^28,29^: 1)haplotype-dominant gene pairs (Hap), where one allele exhibits higher expression in at least one-third of the tissues compared to its partner allele, with no significant difference in other tissues; 2)sub/neo-functionalized alleles (SubNeo), where both alleles show significantly higher expression in at least one tissue; 3)NoDiff, where no differential expression is observed between the alleles. Allelic bias was evaluated using a mixed-effects linear model from the lme4^94^ package in R, and the minor allele contribution was calculated. LOESS regression was used to assess the correlation between total expression levels and allelic ratio.

### Identification of homeologous and orthologous gene sets

To identify homologous relationships among *I. pseudotinctoria* and other species, we downloaded their protein sequences and aligned them using OrthoFinder^95^. Firstly, protein sets were collected from 16 sequenced species and the longest transcripts of each gene were extracted, in which miscoded genes and genes exhibiting premature termination were discarded. Proteins with no homologs in the other 15 genomes were extracted as species-specific genes including *I. pseudotinctoria* specific unique genes and unclustered genes. Functional annotation of species-specific genes and the enrichment tests were performed using information from homologs in the Gene Ontology ( http://www.geneontology.org/ ) and KEGG ( Kyoto Encyclopedia of Genes and Genomes ) database.

### Phylogenetic analyses and gene family expansion/contraction analysis

On the basis of the identified orthologous gene sets with OrthoFinder^95^, molecular phylogenetic analysis was performed using the shared single-copy genes. Briefly, the coding sequences were extracted from the single-copy families and each ortholog group were multiple aligned using Mafft ^96^. Poorly aligned sequences were then eliminated using Gblocks^97^ and the RAxML-NG^98^ was used for the phylogenetic tree construction with 100 bootstrap replicates. Based on the phylogenetic tree, PAML^99^ was utilized to compute the mean substitution rates along each branch and estimate the species divergent time. Significant expansion or contraction of specific gene families is often associated with adaptive divergence of closely related species. According to the results of OrthoFinder^95^, expansions and contractions of orthologous gene families were then detected using CAFÉ^100^ which uses a birth and death process to model gene gain and loss over a phylogeny.

### Screening for whole genome duplication events

Synonymous substitution rate (Ks) estimation were used to detect whole-genome duplication (WGD) events in the *I. pseudotinctoria* genome. Firstly, protein sequences of *I. pseudotinctoria* were extracted and all-vs-all paralog analysis were performed using best hits from primary protein sequences by self-BLASTp in these plants. After filtering by identity and coverage, the BLASTP^101^ results were then subjected to MCScanX ^102^ and the respective collinear blocks were thus identified. Finally, the Ks were then calculated for the syntenic blocks gene pairs using KaKs_Calculator^103^ and potential WGD events in each genome were evaluated based on their ks distribution.

### UPLC-MS/MS detection

The UPLC–MS/MS analysis was performed using an ExionLC™ AD UPLC (SCIEX) coupled with a QTRAP 6500+ mass spectrometer for metabolite detection. Plant tissue samples (50 mg, freeze-dried) were extracted with 70% cold methanol. After vortexing and ultrasonication, the samples were centrifuged at 12,000 rpm for 3 minutes, and the supernatant was filtered through a 0.22 μm filter before injection. Chromatographic separation was achieved using an Agilent SB-C18 column (1.8 μm, 2.1 × 100 mm), with mobile phases A: 0.1% formic acid in water and B: 0.1% formic acid in acetonitrile. The gradient program was 0–9 min: B 5%→95%, held for 1 min, and then 10.00–11.10 min back to 5%, equilibrated to 14 min. The flow rate was 0.35 mL/min, and the column was maintained at 40°C, with an injection volume of 2 μL.

Mass spectrometry was performed using electrospray ionization (ESI) in both positive and negative modes, with the following parameters: ion source temperature 500°C, ion spray voltage +5500 V / −4500 V, and the gas pressures GSI = 50 psi, GSII = 60 psi, CUR = 25 psi, with medium collision gas. Data were acquired in MRM mode. Metabolite identification was based on retention time, Q1/Q3 ion pairs, and fragmentation patterns compared to standards and database entries. The data were processed using Analyst 1.6.3 and MultiQuant for peak detection, integration, and quantification. A QC sample was inserted for every 10 samples to monitor system stability (TIC/XIC overlay). Data preprocessing involved filling missing values with the 1/5 of the minimum value for each metabolite and retaining metabolites with QC CV < 0.5 for analysis.

### Transcriptional regulation of flavonoid biosynthesis

We conducted a comprehensive gene expression and co-expression network analysis using the entire transcriptome dataset to uncover the transcriptional regulatory network involving flavonoid biosynthesis genes and transcription factors (TFs). Genes with TPM < 1 across all tissues were filtered out, and the remaining genes were used to construct a co-expression network via WGCNA^104^. The co-expression modules were identified using the block module function with the following parameters: soft threshold power = 16; TOMtype = signed; mergeCutHeight = 0.25; and minModuleSize = 50. Transcription factors (TFs) in the *I. pseudotinctoria* genome were then identified using PlantTFDB^105^ with default parameters. Finally, the gene-TF interaction network was visualized in Cytoscape^106^.

## Acknowledgments

This work was supported by funds from Livestock Science and Technology Innovation Team Cultivation Project (22535C to Y.F), Chongqing performance incentive guide special project (CSTB2023JXJL-YFX0034 to Y.F), Chongqing Modern Agricultural Industry Technology System (CQMAITS202613 to Y.F), Chongqing Technology Innovation and Application Development Project (CSTB2023TIAD-KPX0024 to Y.F), National Natural Science Foundation of China (32271753 & 32471764 to X.M.), the National Key R&D Program of China (2024YFD1301202 to X.M.). Some of the analytical work was completed on the High-performance Computing Platform of Sichuan Agricultural University.

## Author contributions

J.P., JM.Z., X.M., and Y.F. designed the research. J.P., J.Z., W.H., Y.X., Y.Xu., C.L., Q.R., and J.C. performed the experiments and generated data J.P., JM.Z., Y.X., and Y.Xu. analyzed the data. J.P. and X.M. wrote the manuscript. All authors reviewed and edited the manuscript.

## Code availability

Customized code and scripts supporting this work are available via GitHub at https://github.com/Jinghanpeng313/Indigofera-pseudotinctoria-T2T-genome.git.

## Reference

1. Du Preez, B. et al. Global biogeographic patterns of the genus Indigofera (Fabaceae: Indigofereae). Braz. J. Bot 48, 19 (2025).

2. du Preez, B. et al. Phylogeny and new sectional classification for the Cape Clade of the genus Indigofera (Fabaceae: Indigofereae). TAXON 74, 819–856 (2025).

3. Rajani, V., Umadevi, S. & Naga Raju, C. A Review on Exploring the Phytochemical and Pharmacological Significance of Indigofera astragalina. Pharmacognosy Magazine 20, 363–371 (2024).

4. Indigofera pseudotinctoria Matsum. | Plants of the World Online | Kew Science. Plants of the World Online http://powo.science.kew.org/taxon/urn:lsid:ipni.org:names:499871-1.

5. Zhao, J. et al. Organelle genomes of Indigofera amblyantha and Indigofera pseudotinctoria: comparative genome analysis, and intracellular gene transfer. Industrial Crops and Products 198, 116674 (2023).

6. Chen, J. et al. Genome Survey Sequencing of Indigofera pseudotinctoria and Identification of Its SSR Markers. Genes 16, 991 (2025).

7. Wen, E. & Liang, H. [Chemical constituents of Indigofera pseudotinctoria]. Zhongguo Zhong Yao Za Zhi 35, 2708–2711 (2010).

8. Rahman, T. U. et al. Phytochemistry and Pharmacology of Genus Indigofera: A Review. Rec. Nat. Prod. 12, 1–13 (2017).

9. Bakasso, S. et al. Polyphenol contents and antioxidant activities of five Indigofera species (Fabaceae) from Burkina Faso. Pak J Biol Sci 11, 1429–1435 (2008).

10. Zhou Jing. The studies on theanti-inflammatory chemical constitutes from Indigofera pseudotinctoria Mats. (Huazhong University, 2012).

11. Wen, E. & Liang, H. Chemical constituents of Indigofera pseudotinctoria. China Journal of Chinese Materia Medica 35, 2708–2711 (2010).

12. B. Y, M., Rm, A., Mo, E. & Babandi, A. Anti-oxidant, Anti-inflammatory, Anti-proliferative and Anti-microbial activities (In vitro) of Indigofera hirsuta and Afrormosia laxiflora. SJMPS 05, 923–930 (2019).

13. Fan, Y. et al. Analysis of Genetic Diversity and Structure Pattern of Indigofera Pseudotinctoria in Karst Habitats of the Wushan Mountains Using AFLP Markers. Molecules 22, 1734 (2017).

14. Otao, T., Kobayashi, T. & Uehara, K. Development and characterization of 14 microsatellite markers for Indigofera pseudotinctoria (Fabaceae). Applications in Plant Sciences 4, 1500110 (2016).

15. Yu, B.-D. & Shim, S.-R. The Optimal Seeding Quantity of Lespedeza cyrtobotrya Miquel and Indigofera pseudo-tinctoria MATSUMURA as Leguminous Woody Plants for the Cut-slope Revegetation. Journal of the Korean Society of Environmental Restoration Technology 19, 61–71 (2016).

16. Ma, S. et al. Physiological response and transcriptome analyses of leguminous Indigofera bungeana Walp. to drought stress. PeerJ 11, e15440 (2023).

17. Yu, B.-D. & Shim, S.-R. The Optimal Seeding Quantity of Lespedeza cyrtobotrya Miquel and Indigofera pseudo-tinctoria MATSUMURA as Leguminous Woody Plants for the Cut-slope Revegetation. Journal of the Korean Society of Environmental Restoration Technology 19, 61–71 (2016).

18. Ravelombola, W. et al. Current status of the genetic and agronomic of industrial indigo Indigofera sp. Euphytica 219, 128 (2023).

19. Yang, Q. & Wang, G. Isoflavonoid metabolism in leguminous plants: an update and perspectives. Front. Plant Sci. 15, (2024).

20. Xiaoxia, S., Shufang, M., Xiaoli, S. & Dianxing, W. Brief report on development of indigofera pseudotinctoria mats high flavonoid mutant and anti-oxidation of its exacts. Journal of Nuclear Agricultural Sciences 24, (2010).

21. Li, J., Yu, Q., Liu, C., Zhang, N. & Xu, W. Flavonoids as key players in cold tolerance: molecular insights and applications in horticultural crops. Hortic Res 12, uhae366 (2025).

22. Xu, Y. et al. Engineering plant hosts for high-efficiency accumulation of flavonoids: Advances, challenges and perspectives. Biotechnology Advances 84, 108692 (2025).

23. Gerometta, E., Grondin, I., Smadja, J., Frederich, M. & Gauvin-Bialecki, A. A review of traditional uses, phytochemistry and pharmacology of the genus Indigofera. Journal of Ethnopharmacology 253, 112608 (2020).

24. Méteignier, L.-V., Nützmann, H.-W., Papon, N., Osbourn, A. & Courdavault, V. Emerging mechanistic insights into the regulation of specialized metabolism in plants. Nat. Plants 9, 22–30 (2023).

25. Cheng, H., Concepcion, G. T., Feng, X., Zhang, H. & Li, H. Haplotype-resolved de novo assembly using phased assembly graphs with HiFiasm. Nat Methods 18, 170–175 (2021).

26. Cosentino, R. O., Brink, B. G. & Siegel, T. N. Allele-specific assembly of a eukaryotic genome corrects apparent framesHiFits and reveals a lack of nonsense-mediated mRNA decay. NAR Genom Bioinform 3, lqab082 (2021).

27. Zou, C. et al. A multitiered haplotype strategy to enhance phased assembly and fine mapping of a disease resistance locus. Plant Physiol 193, 2321–2336 (2023).

28. Zhang, X. et al. Haplotype-resolved genome assembly provides insights into evolutionary history of the tea plant Camellia sinensis. Nat Genet 53, 1250–1259 (2021).

29. Sun, M. et al. Haplotype-resolved, gap-free genome assemblies provide insights into the divergence between Asian and European pears. Nat Genet 57, 2040–2051 (2025).

30. Li, C. et al. The haplotype-resolved telomere-to-telomere genome and OMICS analyses reveal genetic responses to tapping in rubber tree. Nat Commun 16, 6255 (2025).

31. Shi, T.-L. et al. High-quality genome assembly enables prediction of allele-specific gene expression in hybrid poplar. Plant Physiol 195, 652–670 (2024).

32. Cheng, H., Concepcion, G. T., Feng, X., Zhang, H. & Li, H. Haplotype-resolved de novo assembly using phased assembly graphs with HiFiasm. Nat Methods 18, 170–175 (2021).

33. Nurk, S. et al. HiCanu: accurate assembly of segmental duplications, satellites, and allelic variants from high-fidelity long reads. Genome Res 30, 1291–1305 (2020).

34. Hu, J. et al. NextDenovo: an efficient error correction and accurate assembly tool for noisy long reads. Genome Biology 25, 107 (2024).

35. Ni, L. et al. Pan-3D genome analysis reveals structural and functional differentiation of soybean genomes. Genome Biology 24, 12 (2023).

36. Wang, L. et al. Pangenome analysis provides insights into legume evolution and breeding. Nat Genet 57, 2052–2061 (2025).

37. Hu, W. et al. Allele-defined genome reveals biallelic differentiation during cassava evolution. Mol Plant 14, 851–854 (2021).

38. Wang, Y. et al. Comparative genomic analyses reveal different genetic basis of two types of fruit in Maloideae. Nat Commun 16, 7463 (2025).

39. Sun, X. et al. Phased diploid genome assemblies and pan-genomes provide insights into the genetic history of apple domestication. Nat Genet 52, 1423–1432 (2020).

40. Zhao, Y. et al. Nuclear phylotranscriptomics and phylogenomics support numerous polyploidization events and hypotheses for the evolution of rhizobial nitrogen-fixing symbiosis in Fabaceae. Molecular Plant 14, 748–773 (2021).

41. Hasan, A., Farman, M. & Ahmed, I. Flavonoid glycosides from Indigofera hebepetala. Phytochemistry 35, 275–276 (1993).

42. Ahmed, H. S. Neuropharmacological effects of calycosin: a translational review of molecular mechanisms and therapeutic applications. Naunyn Schmiedebergs Arch Pharmacol 398, 12891–12909 (2025).

43. Tuli, H. S. et al. Molecular mechanisms underlying chemopreventive potential of butein: Current trends and future perspectives. Chem Biol Interact 350, 109699 (2021).

44. Aboulaghras, S. et al. Health Benefits and Pharmacological Aspects of Chrysoeriol. Pharmaceuticals (Basel) 15, 973 (2022).

45. Hichri, I. et al. Recent advances in the transcriptional regulation of the flavonoid biosynthetic pathway. J Exp Bot 62, 2465–2483 (2011).

46. Patil, J. R. et al. Flavonoids in plant-environment interactions and stress responses. Discov. Plants 1, 68 (2024).

47. Kuljarusnont, S., Iwakami, S., Iwashina, T. & Tungmunnithum, D. Flavonoids and Other Phenolic Compounds for Physiological Roles, Plant Species Delimitation, and Medical Benefits: A Promising View. Molecules 29, 5351 (2024).

48. Li, P. et al. Diverse roles of MYB transcription factors in regulating secondary metabolite biosynthesis, shoot development, and stress responses in tea plants (Camellia sinensis). The Plant Journal 110, 1144–1165 (2022).

49. Zhang, D. et al. Diverse roles of MYB transcription factors in plants. Journal of Integrative Plant Biology 67, 539–562 (2025).

50. Wang, F. et al. AtMYB12 regulates flavonoids accumulation and abiotic stress tolerance in transgenic Arabidopsis thaliana. Mol Genet Genomics 291, 1545–1559 (2016).

51. Chu, S. et al. An R2R3-type MYB transcription factor, GmMYB29, regulates isoflavone biosynthesis in soybean. PLOS Genetics 13, e1006770 (2017).

52. Rahman, P. & Mehnaz, S. International Journal for Multidisciplinary Research (IJFMR). SSRN Journal https://doi.org/10.2139/ssrn.5054029 (2024) doi:10.2139/ssrn.5054029.

53. Zhao, J. Flavonoid transport mechanisms: how to go, and with whom. Trends Plant Sci 20, 576–585 (2015).

54. Buer, C. S., Muday, G. K. & Djordjevic, M. A. Flavonoids are differentially taken up and transported long distances in Arabidopsis. Plant Physiol 145, 478–490 (2007).

55. Biala, W., Banasiak, J., Jarzyniak, K., Pawela, A. & Jasinski, M. Medicago truncatula ABCG10 is a transporter of 4-coumarate and liquiritigenin in the medicarpin biosynthetic pathway. J Exp Bot 68, 3231–3241 (2017).

56. Ng, M.-S. et al. MATE-Type Proteins Are Responsible for Isoflavone Transportation and Accumulation in Soybean Seeds. International Journal of Molecular Sciences 22, 12017 (2021).

57. Yu, S. et al. Dissection of the spatial dynamics of biosynthesis, transport, and turnover of major amino acids in tea plants (Camellia sinensis). Hortic Res 11, uhae060 (2024).

58. Gerometta, E., Grondin, I., Smadja, J., Frederich, M. & Gauvin-Bialecki, A. A review of traditional uses, phytochemistry and pharmacology of the genus Indigofera. Journal of Ethnopharmacology 253, 112608 (2020).

59. Biała, W., Banasiak, J., Jarzyniak, K., Pawela, A. & Jasiński, M. Medicago truncatula ABCG10 is a transporter of 4-coumarate and liquiritigenin in the medicarpin biosynthetic pathway. J Exp Bot 68, 3231–3241 (2017).

60. Zeng, X. et al. Chromosome-level scaffolding of haplotype-resolved assemblies using Hi-C data without reference genomes. Nat. Plants 10, 1184–1200 (2024).

61. Robinson, J. T. et al. Juicebox.js Provides a Cloud-Based Visualization System for Hi-C Data. Cell Syst 6, 256–258.e1 (2018).

62. Xu, M. et al. TGS-GapCloser: A fast and accurate gap closer for large genomes with low coverage of error-prone long reads. Gigascience 9, giaa094 (2020).

63. Lin, Y. et al. quarTeT: a telomere-to-telomere toolkit for gap-free genome assembly and centromeric repeat identification. Hortic Res 10, uhad127 (2023).

64. Xu, D. et al. CentIER: Accurate centromere identification for plant genomes. Plant Commun 5, 101046 (2024).

65. Ellinghaus, D., Kurtz, S. & Willhoeft, U. LTRharvest, an efficient and flexible software for de novo detection of LTR retrotransposons. BMC Bioinformatics 9, 18 (2008).

66. Xu, Z. & Wang, H. LTR_FINDER: an efficient tool for the prediction of full-length LTR retrotransposons. Nucleic Acids Research 35, W265–W268 (2007).

67. Ou, S. & Jiang, N. LTR_retriever: A Highly Accurate and Sensitive Program for Identification of Long Terminal Repeat Retrotransposons. Plant Physiol 176, 1410–1422 (2018).

68. Flynn, J. M. et al. RepeatModeler2 for automated genomic discovery of transposable element families. Proc Natl Acad Sci U S A 117, 9451–9457 (2020).

69. Tarailo-Graovac, M. & Chen, N. Using RepeatMasker to identify repetitive elements in genomic sequences. Curr Protoc Bioinformatics Chapter 4, 4.10.1–4.10.14 (2009).

70. Storer, J., Hubley, R., Rosen, J., Wheeler, T. J. & Smit, A. F. The Dfam community resource of transposable element families, sequence models, and genome annotations. Mobile DNA 12, 2 (2021).

71. Jurka, J. et al. Repbase Update, a database of eukaryotic repetitive elements. Cytogenet Genome Res 110, 462–467 (2005).

72. Campbell, M. S. et al. MAKER-P: A Tool Kit for the Rapid Creation, Management, and Quality Control of Plant Genome Annotations. Plant Physiol 164, 513–524 (2014).

73. Rognes, T., Flouri, T., Nichols, B., Quince, C. & Mahé, F. VSEARCH: a versatile open source tool for metagenomics. PeerJ 4, e2584 (2016).

74. Yan, H., Bombarely, A. & Li, S. DeepTE: a computational method for de novo classification of transposons with convolutional neural network. Bioinformatics 36, 4269–4275 (2020).

75. Benson, G. Tandem repeats finder: a program to analyze DNA sequences. Nucleic Acids Res 27, 573–580 (1999).

76. Tarailo-Graovac, M. & Chen, N. Using RepeatMasker to Identify Repetitive Elements in Genomic Sequences. Current Protocols in Bioinformatics 25, 4.10.1–4.10.14 (2009).

77. Nawrocki, E. P. & Eddy, S. R. Infernal 1.1: 100-fold faster RNA homology searches. Bioinformatics 29, 2933–2935 (2013).

78. Lagesen, K. et al. RNAmmer: consistent and rapid annotation of ribosomal RNA genes. Nucleic Acids Res 35, 3100–3108 (2007).

79. Chan, P. P., Lin, B. Y., Mak, A. J. & Lowe, T. M. tRNAscan-SE 2.0: improved detection and functional classification of transfer RNA genes. Nucleic Acids Res 49, 9077–9096 (2021).

80. Gabriel, L., et al. BRAKER3: Fully automated genome annotation using RNA-seq and protein evidence with GeneMark-ETP, AUGUSTUS and TSEBRA. bioRxiv 2023.06.10.544449 (2024) doi:10.1101/2023.06.10.544449.

81. Kim, D., Paggi, J. M., Park, C., Bennett, C. & Salzberg, S. L. Graph-based genome alignment and genotyping with HISAT2 and HISAT-genotype. Nat Biotechnol 37, 907–915 (2019).

82. Pertea, M. et al. StringTie enables improved reconstruction of a transcriptome from RNA-seq reads. Nat Biotechnol 33, 290–295 (2015).

83. Keilwagen, J., Hartung, F., Paulini, M., Twardziok, S. O. & Grau, J. Combining RNA-seq data and homology-based gene prediction for plants, animals and fungi. BMC Bioinformatics 19, 189 (2018).

84. Keilwagen, J., Hartung, F. & Grau, J. GeMoMa: Homology-Based Gene Prediction Utilizing Intron Position Conservation and RNA-seq Data. Methods Mol Biol 1962, 161–177 (2019).

85. Marçais, G. et al. MUMmer4: A fast and versatile genome alignment system. PLOS Computational Biology 14, e1005944 (2018).

86. Zhou, Z.-W. et al. GenomeSyn: a bioinformatics tool for visualizing genome synteny and structural variations. Journal of Genetics and Genomics 49, 1174–1176 (2022).

87. Goel, M., Sun, H., Jiao, W.-B. & Schneeberger, K. SyRI: finding genomic rearrangements and local sequence differences from whole-genome assemblies. Genome Biology 20, 277 (2019).

88. Quinlan, A. R. & Hall, I. M. BEDTools: a flexible suite of utilities for comparing genomic features. Bioinformatics 26, 841–842 (2010).

89. Tang, H., et al. JCVI: A versatile toolkit for comparative genomics analysis. iMeta 3, e211 (2024).

90. Buchfink, B., Xie, C. & Huson, D. H. Fast and sensitive protein alignment using DIAMOND. Nat Methods 12, 59–60 (2015).

91. Wu, T. D. & Watanabe, C. K. GMAP: a genomic mapping and alignment program for mRNA and EST sequences. Bioinformatics 21, 1859–1875 (2005).

92. Loman, N. J., Quick, J. & Simpson, J. T. A complete bacterial genome assembled de novo using only nanopore sequencing data. Nat Methods 12, 733–735 (2015).

93. Love, M. I., Huber, W. & Anders, S. Moderated estimation of fold change and dispersion for RNA-seq data with DESeq2. Genome Biology 15, 550 (2014).

94. Bates, D., Mächler, M., Bolker, B. & Walker, S. Fitting Linear Mixed-Effects Models Using lme4. Journal of Statistical Software 67, 1–48 (2015).

95. Emms, D. M. & Kelly, S. OrthoFinder: phylogenetic orthology inference for comparative genomics. Genome Biol 20, 238 (2019).

96. Katoh, K. & Standley, D. M. MAFFT multiple sequence alignment software version 7: improvements in performance and usability. Mol Biol Evol 30, 772–780 (2013).

97. Talavera, G. & Castresana, J. Improvement of Phylogenies after Removing Divergent and Ambiguously Aligned Blocks from Protein Sequence Alignments. Syst Biol 56, 564–577 (2007).

98. Kozlov, A. M., Darriba, D., Flouri, T., Morel, B. & Stamatakis, A. RAxML-NG: a fast, scalable and user-friendly tool for maximum likelihood phylogenetic inference. Bioinformatics 35, 4453–4455 (2019).

99. Yang, Z. PAML 4: phylogenetic analysis by maximum likelihood. Mol Biol Evol 24, 1586–1591 (2007).

100. Mendes, F. K., Vanderpool, D., Fulton, B. & Hahn, M. W. CAFE 5 models variation in evolutionary rates among gene families. Bioinformatics 36, 5516–5518 (2021).

101. Camacho, C., et al. BLAST+: architecture and applications. BMC Bioinformatics 10, 421 (2009).

102. Wang, Y. et al. MCScanX: a toolkit for detection and evolutionary analysis of gene synteny and collinearity. Nucleic Acids Res 40, e49 (2012).

103. Zhang, Z. KaKs_Calculator 3.0: Calculating Selective Pressure on Coding and Non-coding Sequences. Genomics Proteomics Bioinformatics 20, 536–540 (2022).

104. Langfelder, P. & Horvath, S. WGCNA: an R package for weighted correlation network analysis. BMC Bioinformatics 9, 559 (2008).

105. Jin, J. et al. PlantTFDB 4.0: toward a central hub for transcription factors and regulatory interactions in plants. Nucleic Acids Res 45, D1040–D1045 (2017).

106. Shannon, P. et al. Cytoscape: A Software Environment for Integrated Models of Biomolecular Interaction Networks. Genome Res. 13, 2498–2504 (2003).

