## supplemental figures for "A haplotype-resolved T2T genome assembly of Indigofera pseudotinctoria reveals the genetic basis of flavonoid biosynthesis in Chinese Indigo"

### Supplementary Figure

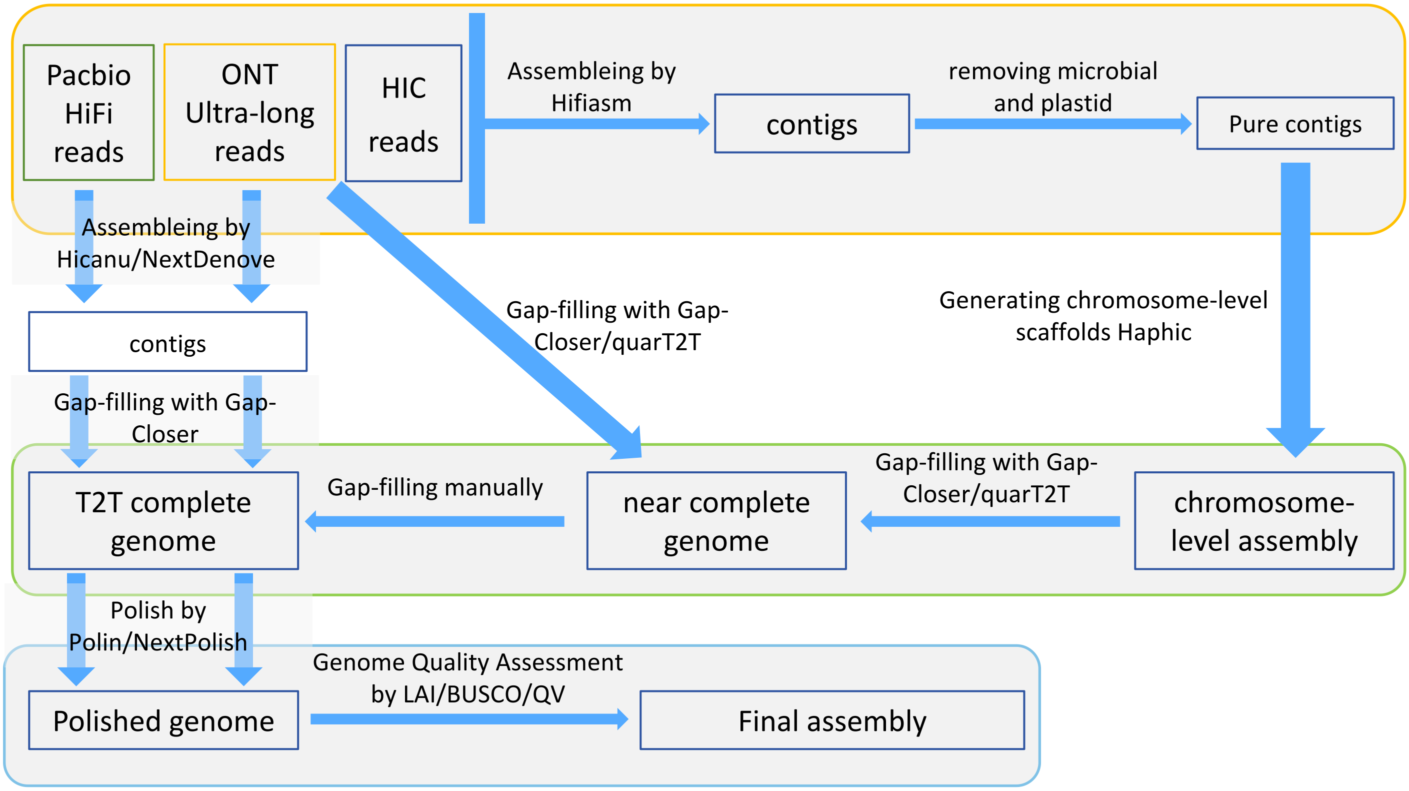


Supplementary Fig.1. Diagram of genome assembly pipeline used in *I. pseudotinctoria* haplotype-resolved genome assembled

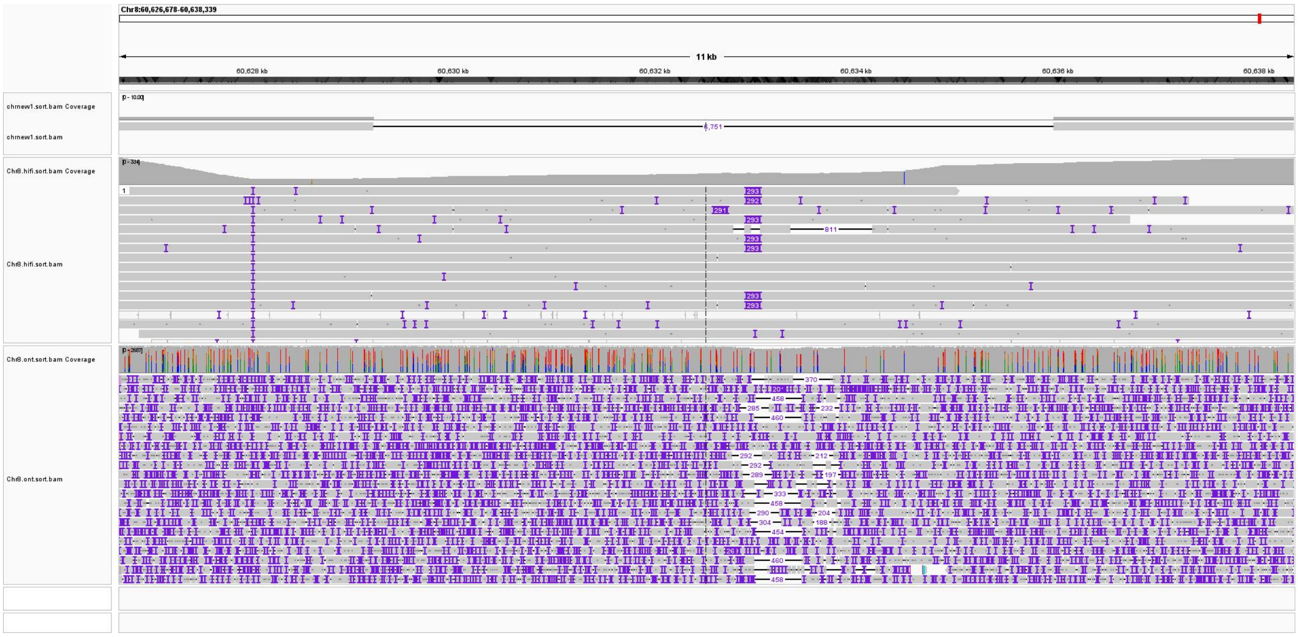


Supplementary Fig.3. An example of closed gaps in Hap2. From top to bottom: filled gap genome, pre-filled gap genome, HiFi reads mapped to filled gap genome, ONT reads filled gap genome.

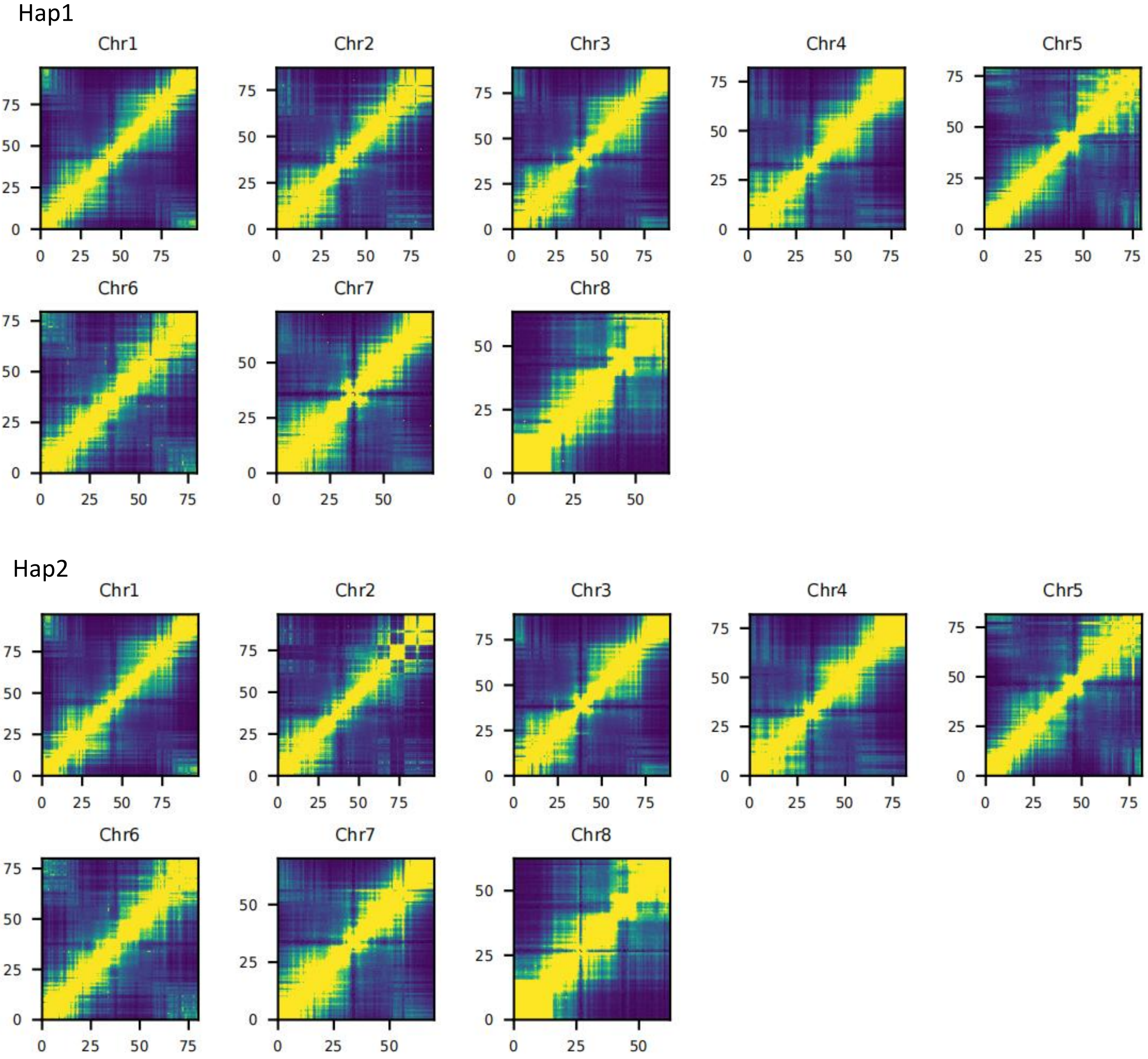


Supplementary Fig.2. Hi-C interaction heatmaps of *I. pseudotinctoria* haplotype-resolved genome.

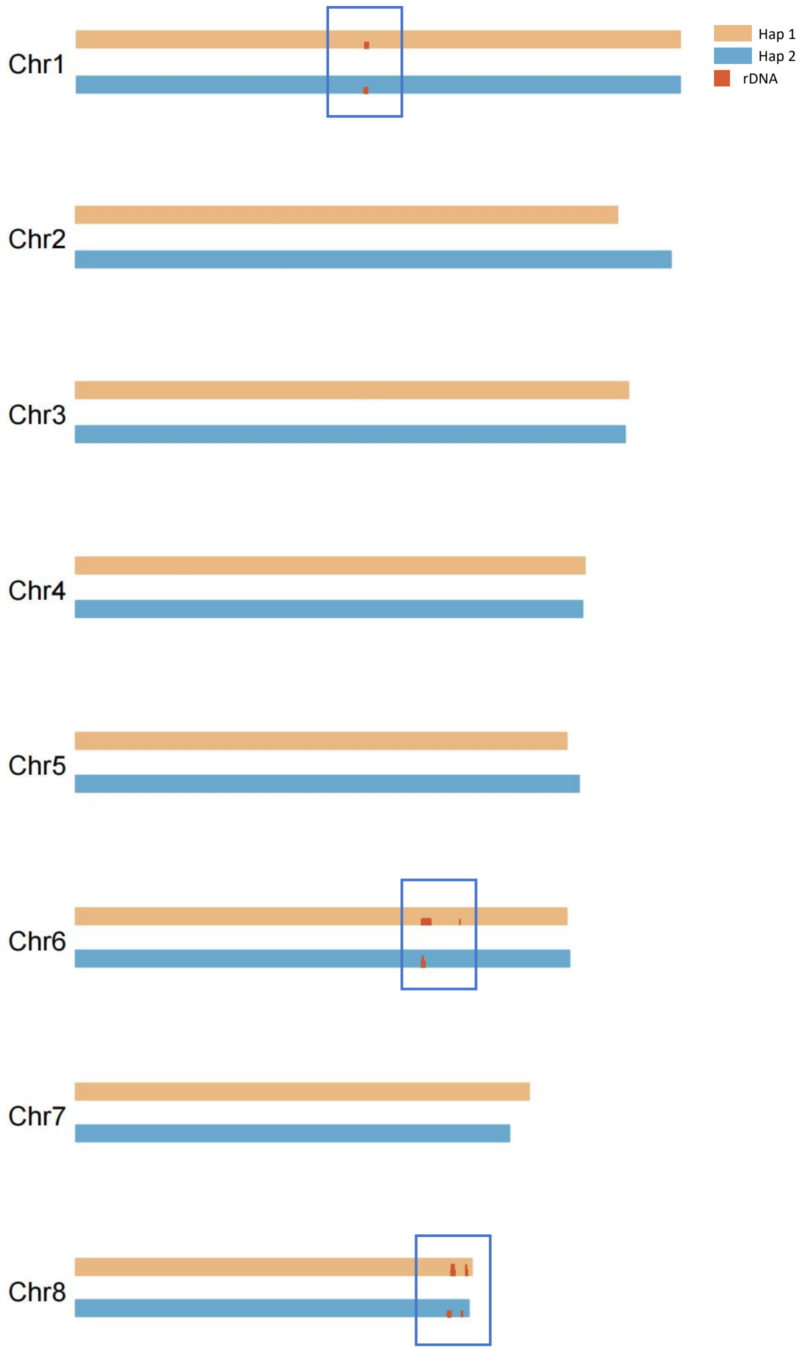


Supplementary Fig.4 Chromosomal alignment of Hap1 and Hap2 with rDNA clusters in *I. pseudotinctoria*.

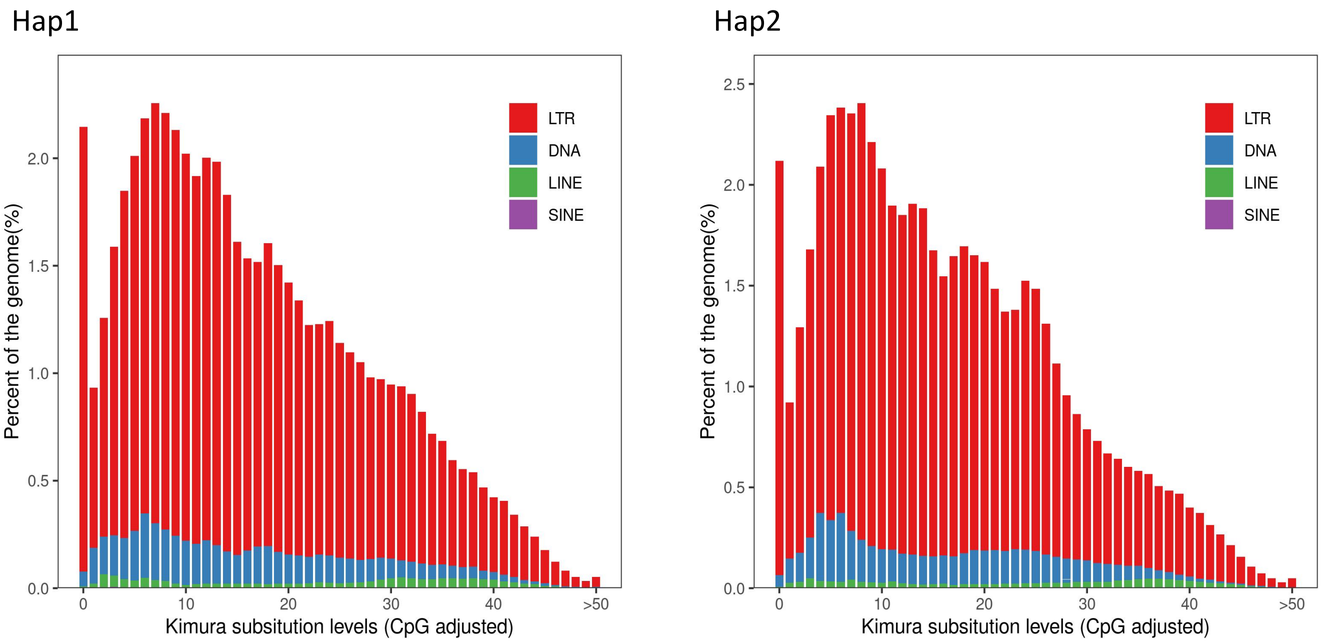


Supplementary Fig.5 Divergence landscape and age distribution of transposable elements (TEs) in the two haplotypes of Indigofera pseudotinctoria.

Hap1 Hap2

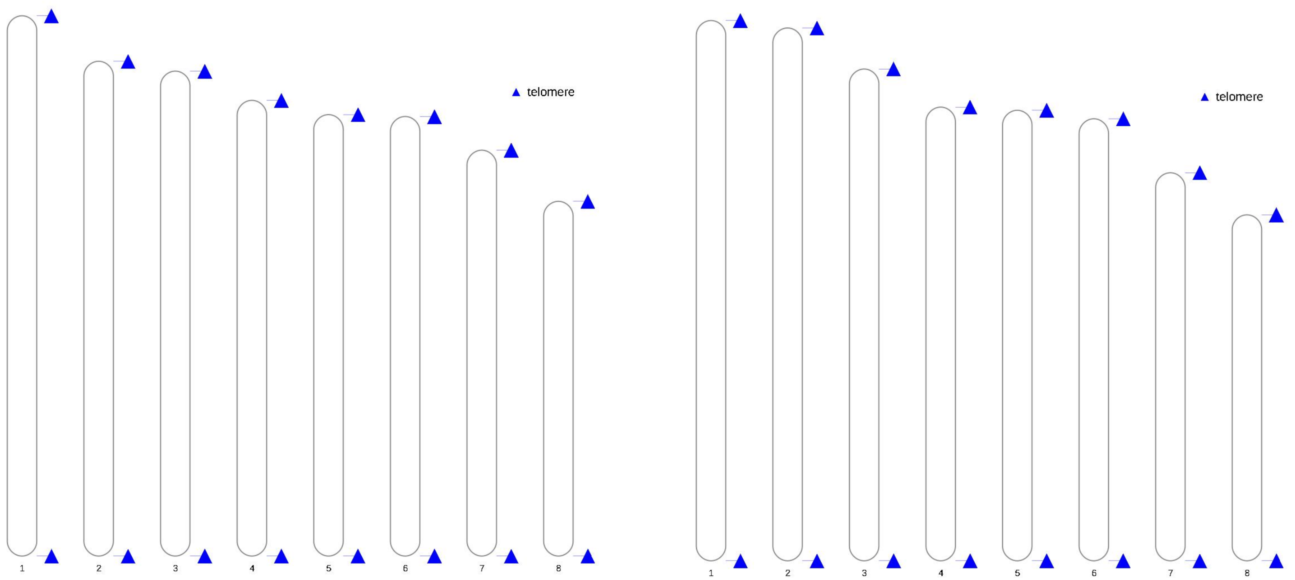


Supplementary Fig.6 Telomeric distribution across the eight chromosomes of the two haplotypes in *I. pseudotinctoria.*

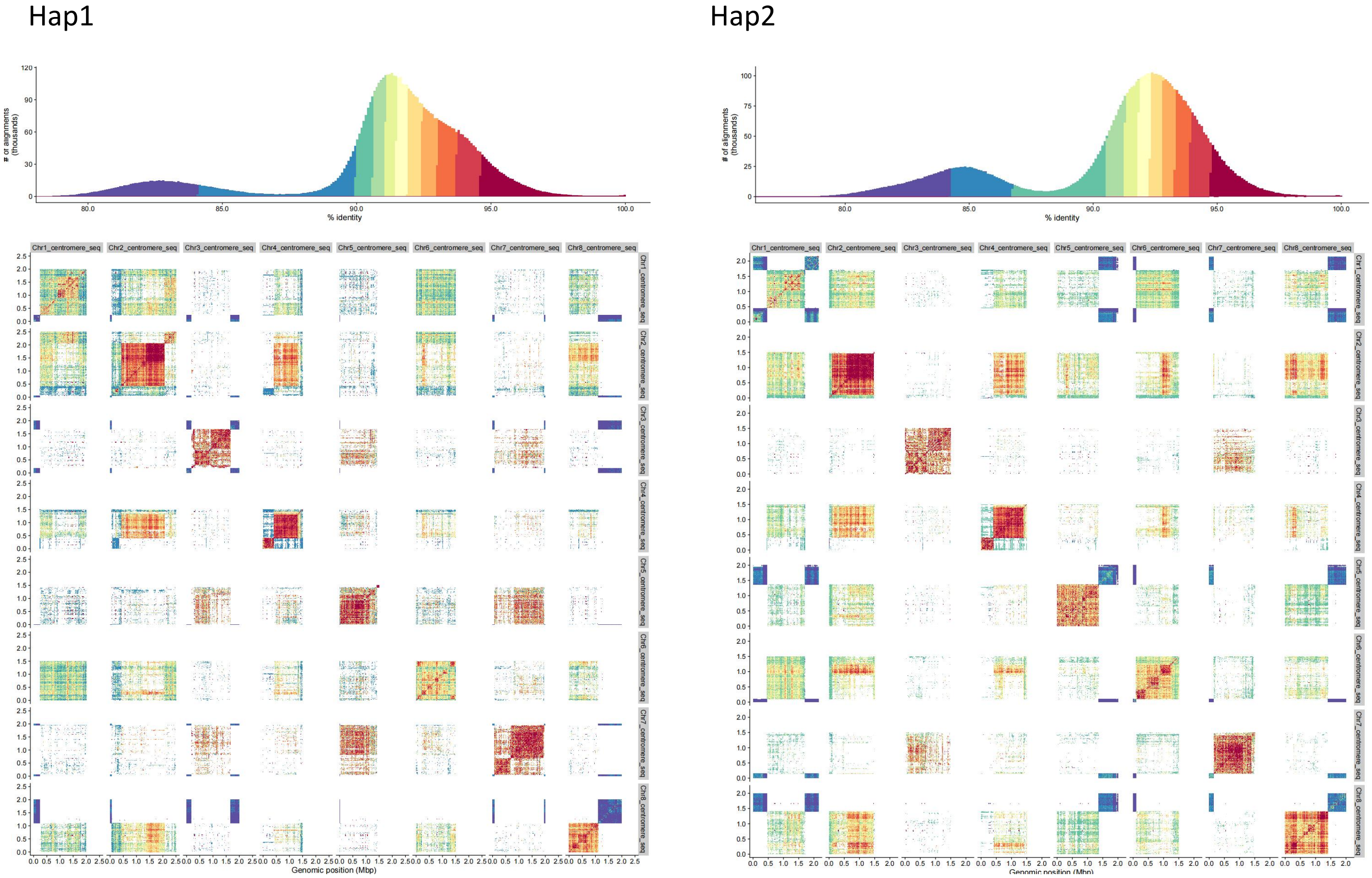


Supplementary Fig. 7. Sequence similarity heatmap of centromeres across chromosomes in *I. pseudotinctoria* haplotype-resolved genome. The sequence similarities among the centromeres were visualized using StainedGlass. The heatmap at the top depicts the number of alignments (Y-axis) with sequence similarity (X-axis, 0-100%) .

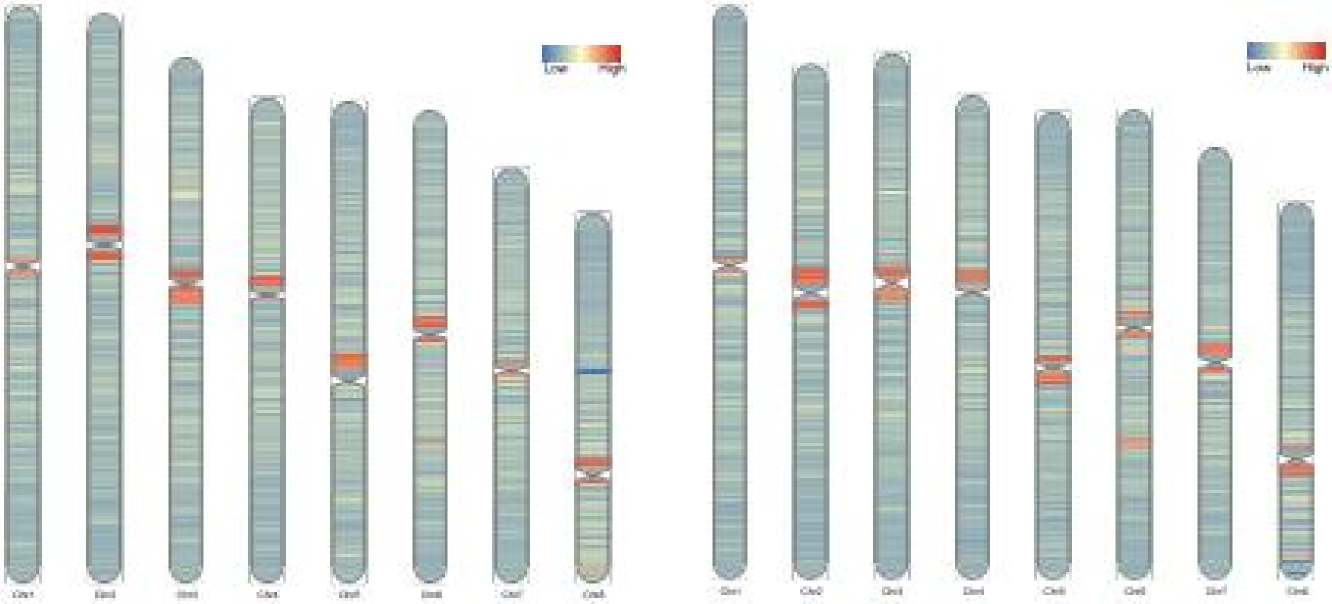


Supplementary Fig.8 DNA methylation landscape confirming centromeric regions in the two haplotypes of *I. pseudotinctoria*.

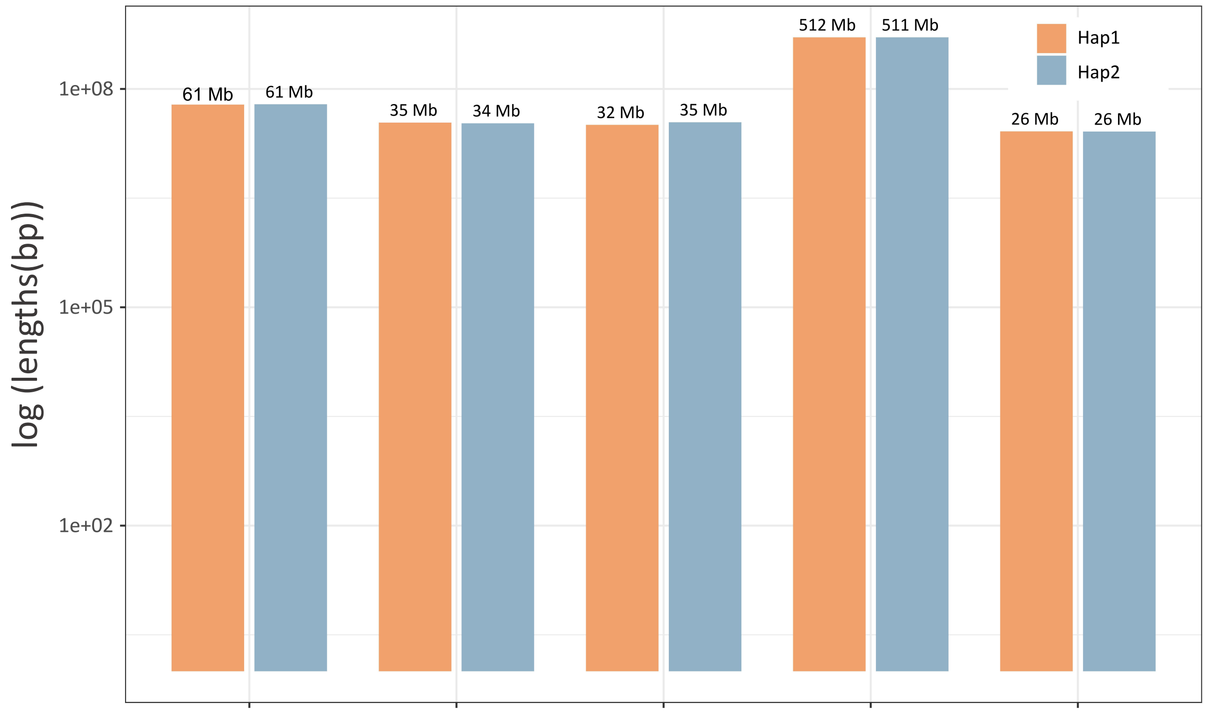


| | Duplication | Inversion | Not aligned | Synteny | Translocation |
|---|---|---|---|---|---|
| Length | 122.21Mb | 70.29Mb | 67.65Mb | 1023.74Mb | 52,64Mb |
| Count | 37,608 | 101 | 18,274 20,280 | 28,969 | 5858 |

Supplementary Fig.9 Statistics on the length and count of structural variations between haplotypes. Length denotes the sum of the lengths of different types of structural variations between two haplotypes. Count indicates the number of different types of structural variations present across the two haplotypes.

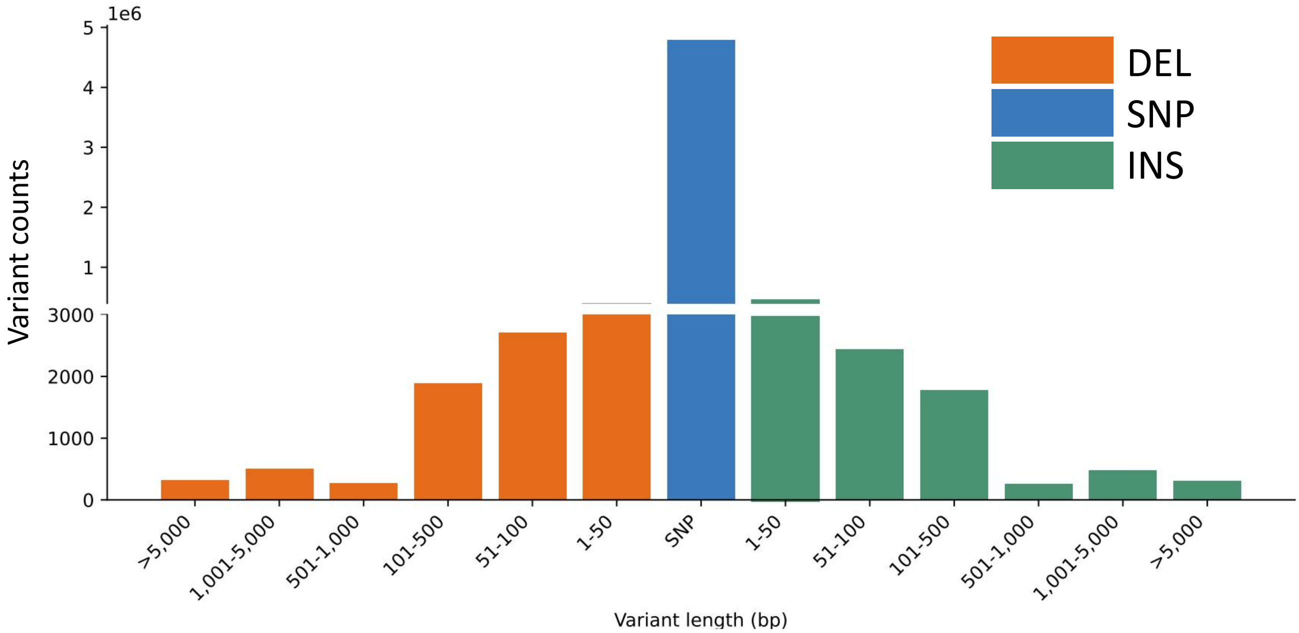


Supplementary Fig.10 Length distribution of DELs, SNPs, and INSs between Hap1 and Hap2 in *I. pseudotinctoria*

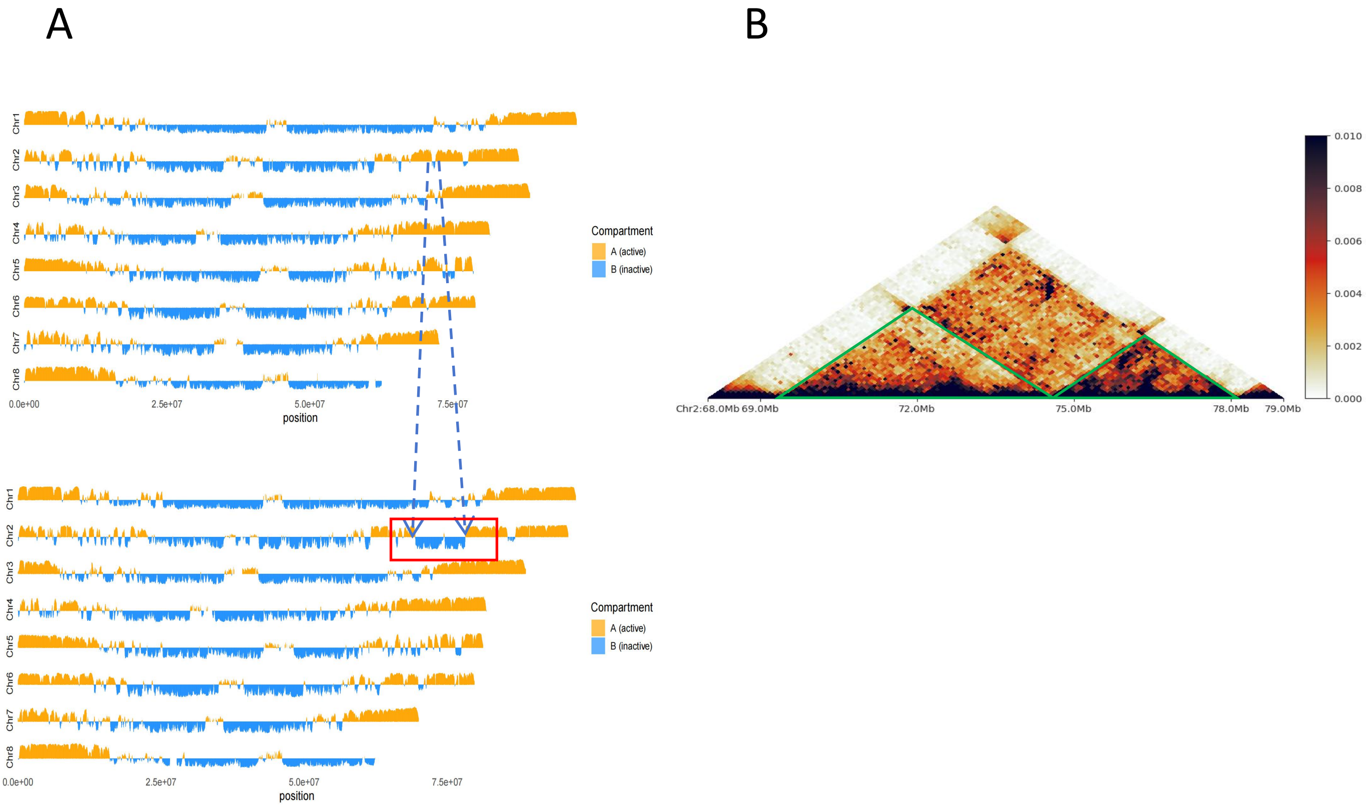


Supplementary Fig.11 Chromatin compartmentalization and Hi-C contact map reveal structural variation between the two haplotypes of *I. pseudotinctoria*. (A) Genome-wide distribution of A/B compartments in Hap1 and Hap2, with active (A, orange) and inactive (B, blue) regions indicated along the eight chromosomes. (B) Hi-C contact heatmap of Chr2 (68–79 Mb) showing the local chromatin topology. The green box marks an independent topologically associating domain (TAD) within the highlighted region.

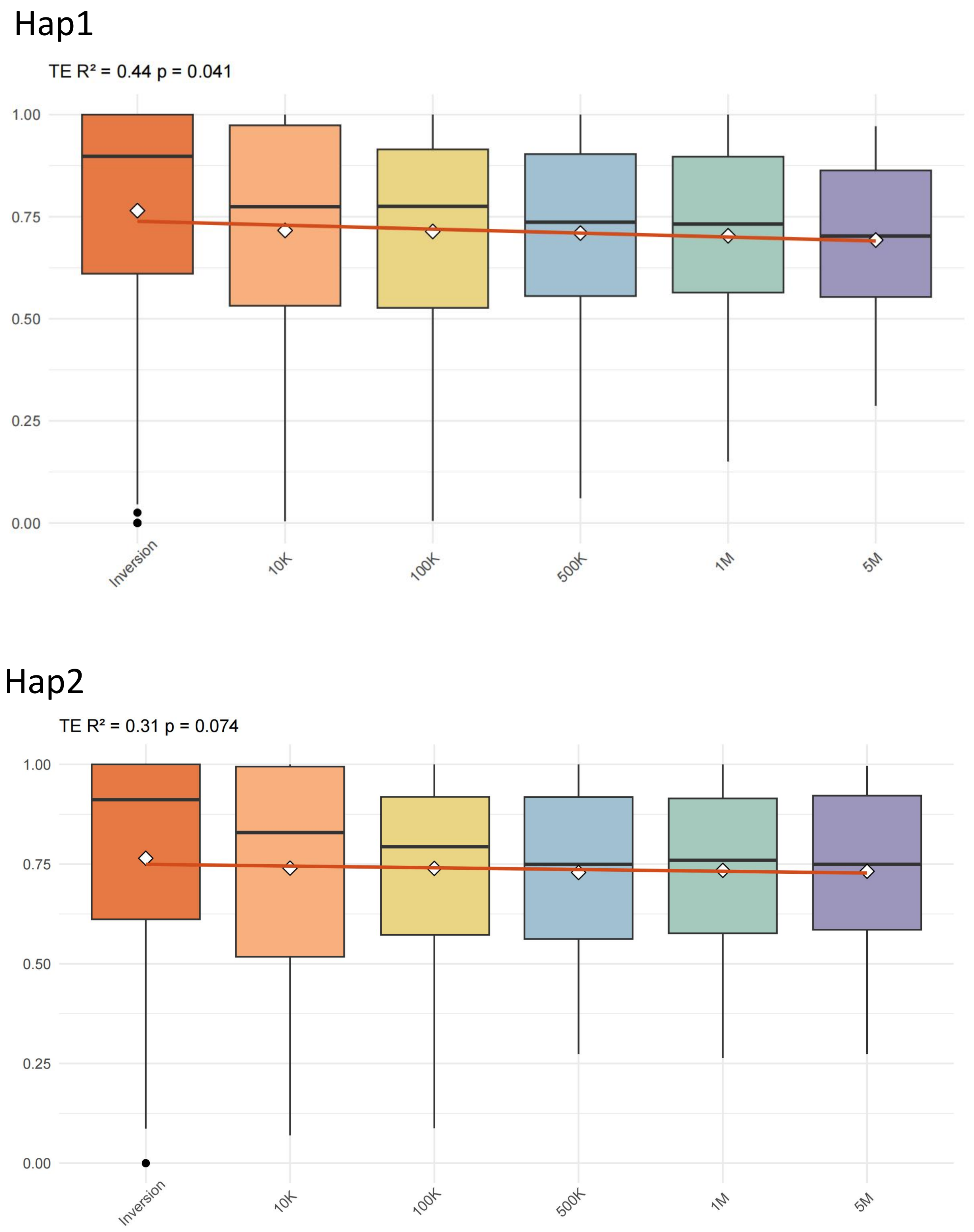


Supplementary Fig.12 Enrichment of transposable elements (TEs) near inversion breakpoints in the two haplotypes of *I. pseudotinctoria*.

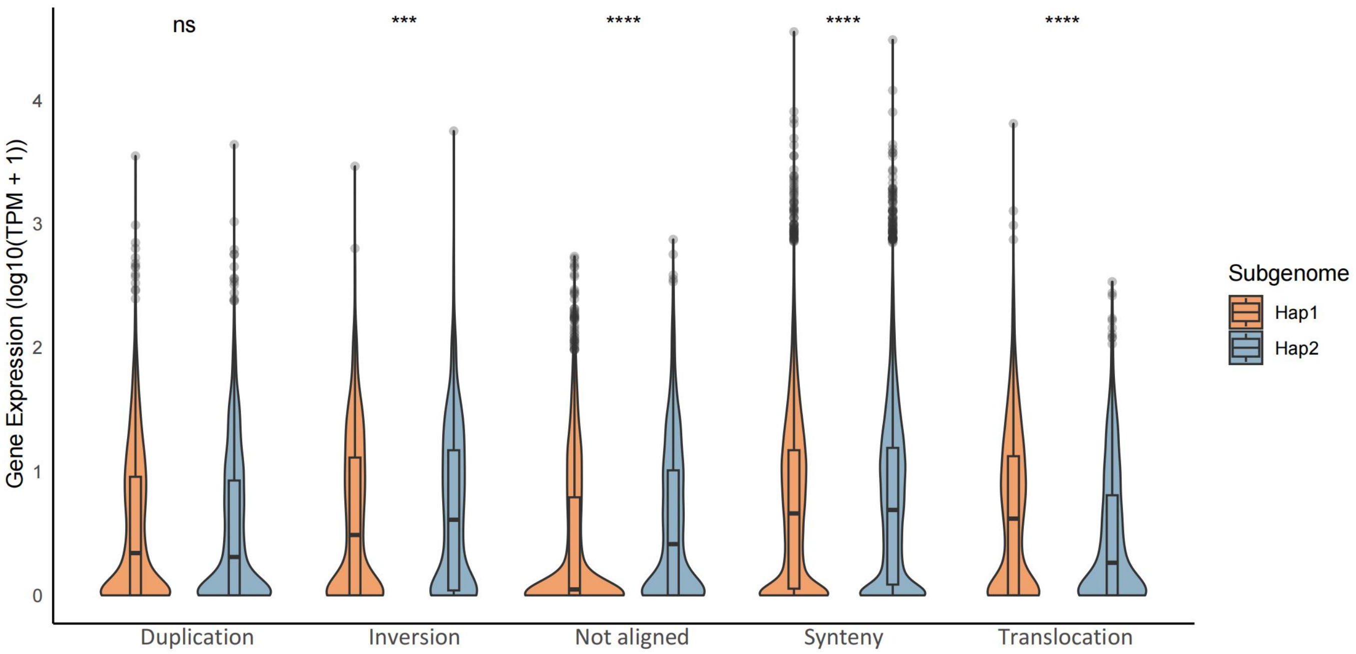


Supplementary Fig.13 Comparison of gene expression between two haplotype genomes across different SV categories. The y-axis represents gene expression levels for genes overlapping with structural variations.

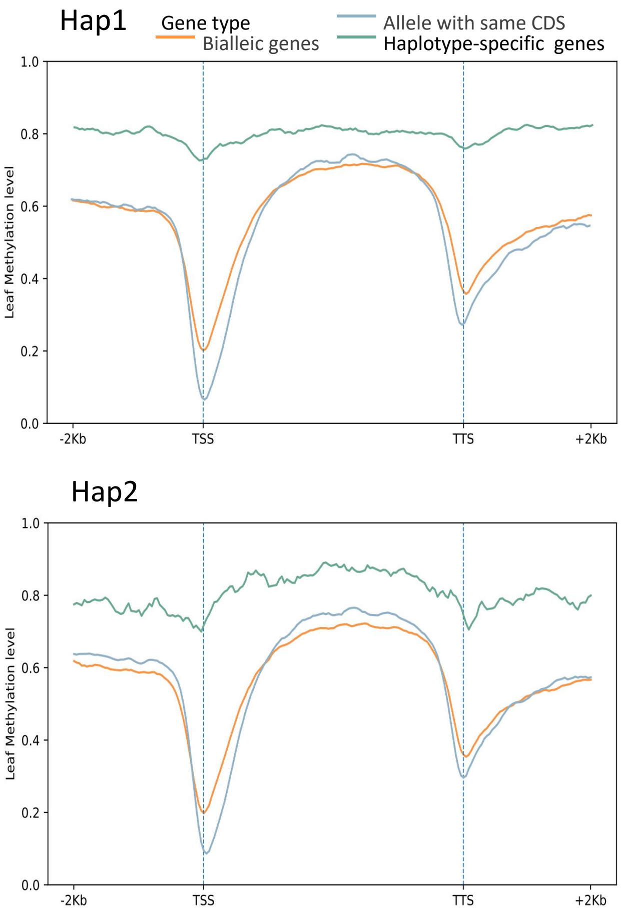


Supplementary Fig.14 DNA methylation profiles of different gene categories around transcriptional start and termination sites in the two haplotypes of *I. pseudotinctoria*.

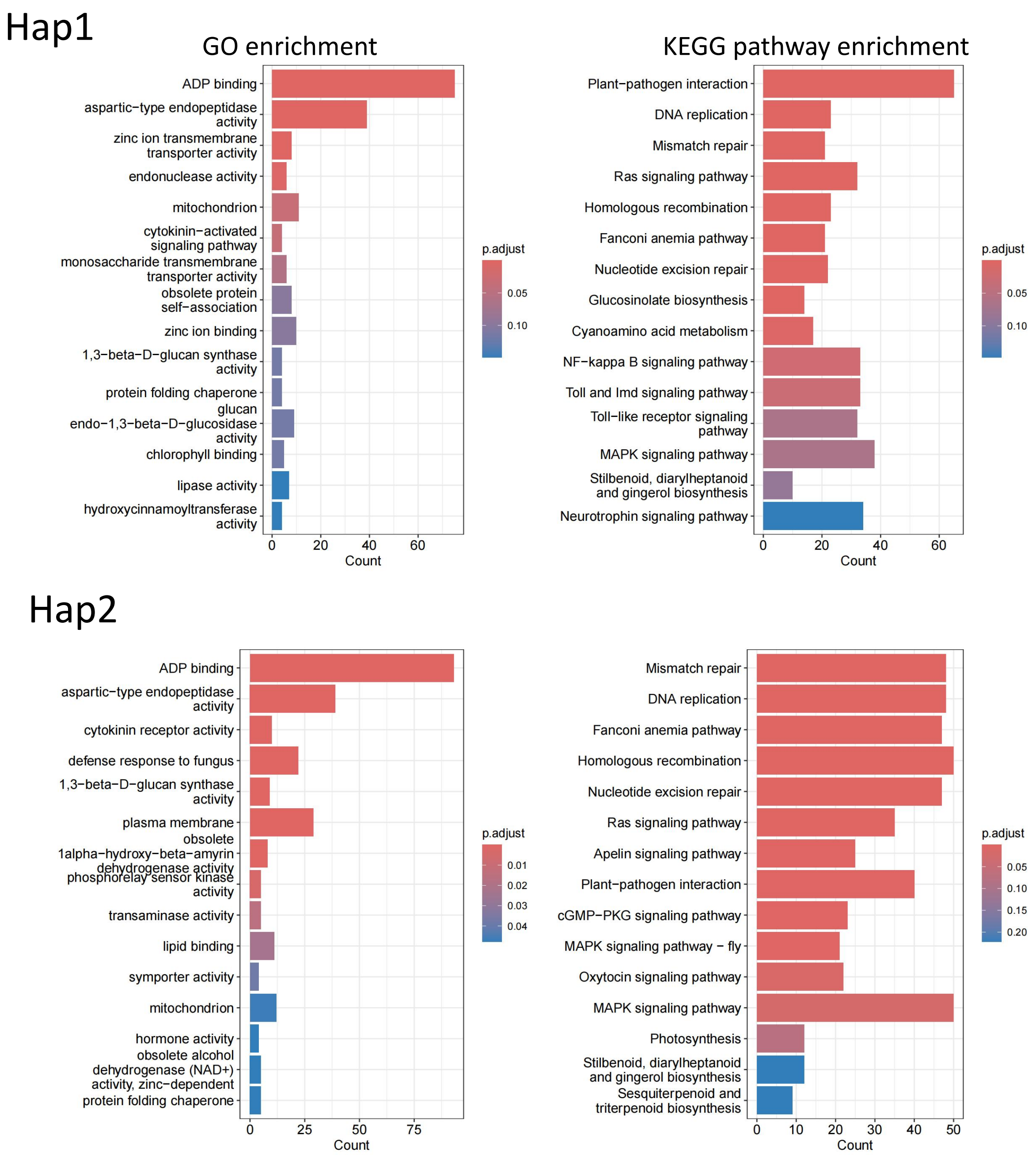


Supplementary Fig.15 GO and KEGG enrichment analyses of haplotype-specific genes in *I. pseudotinctoria*. Functional enrichment of haplotype-specific genes based on Gene Ontology (GO, left panels) and KEGG pathways (right panels) for Hap1 (top) and Hap2 (bottom).

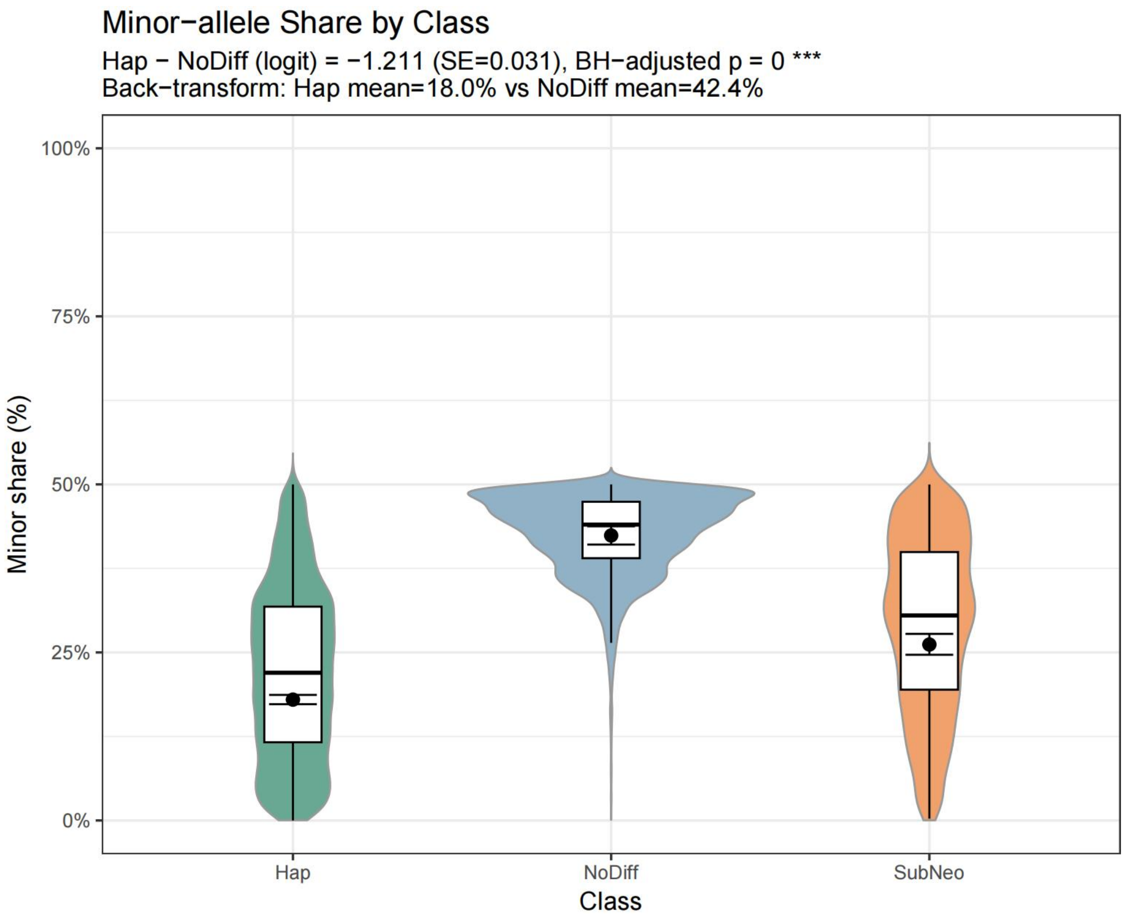


Supplementary Fig.16 Minor-allele share across allelic expression classes in *I. pseudotinctoria*.

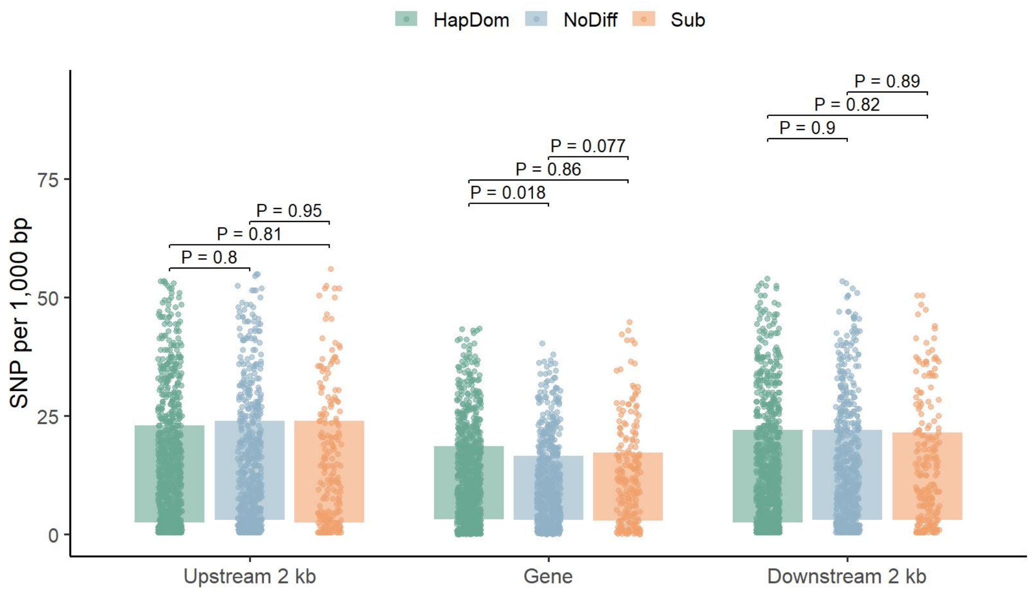


Supplementary Fig.17 The SNP density in gene features of Hap1 in DS (nHapDom = 1252, nSub =307 , nNodiff = 760). The center line is the median, the box limits the first and third quartiles and the whiskers 1.5× the interquartile range (IQR). The terms nHapDom, nSub and nNodiff refer to the counts of different ASE genes. The y axis represents SNP numbers every 1,000 bp. The two-sided Student’s t-test was used for determining the significance.

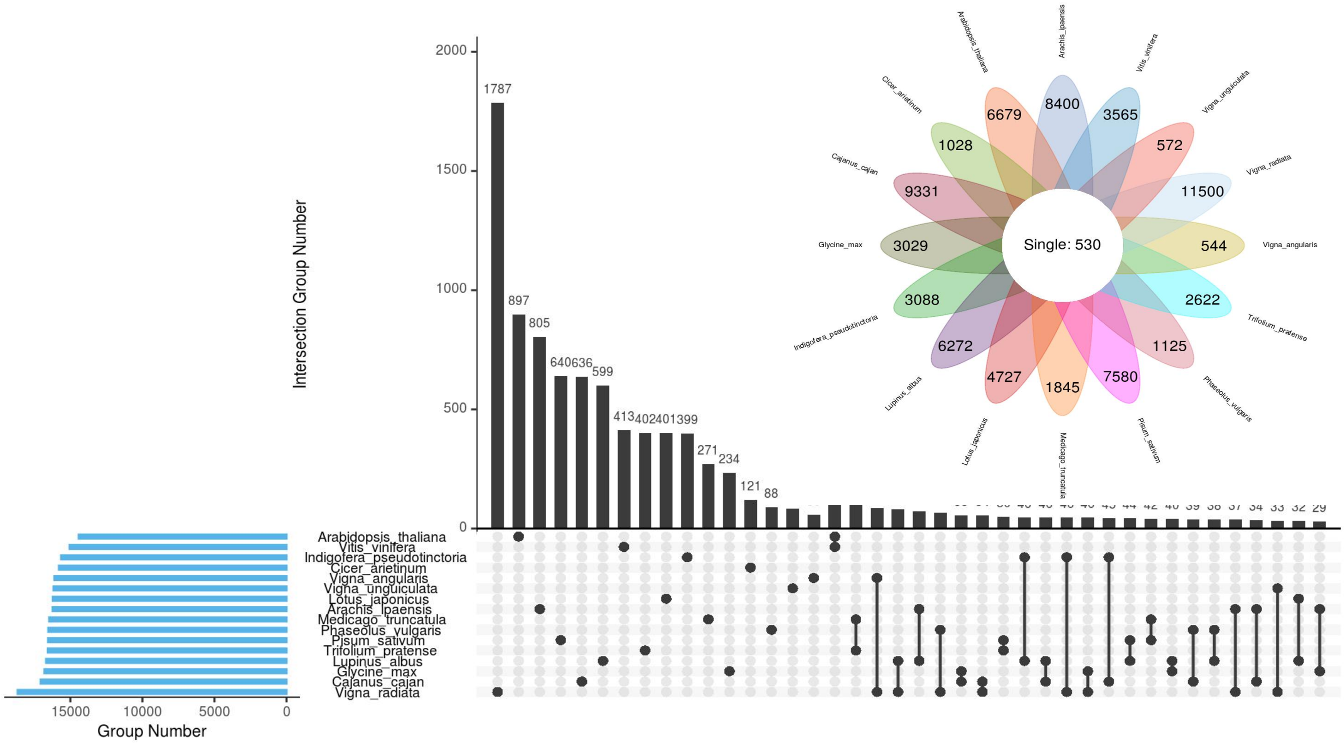


Supplementary Fig.18 The UpSet diagram of homologous genes between species and the petal diagram of genes specific to the species itself.

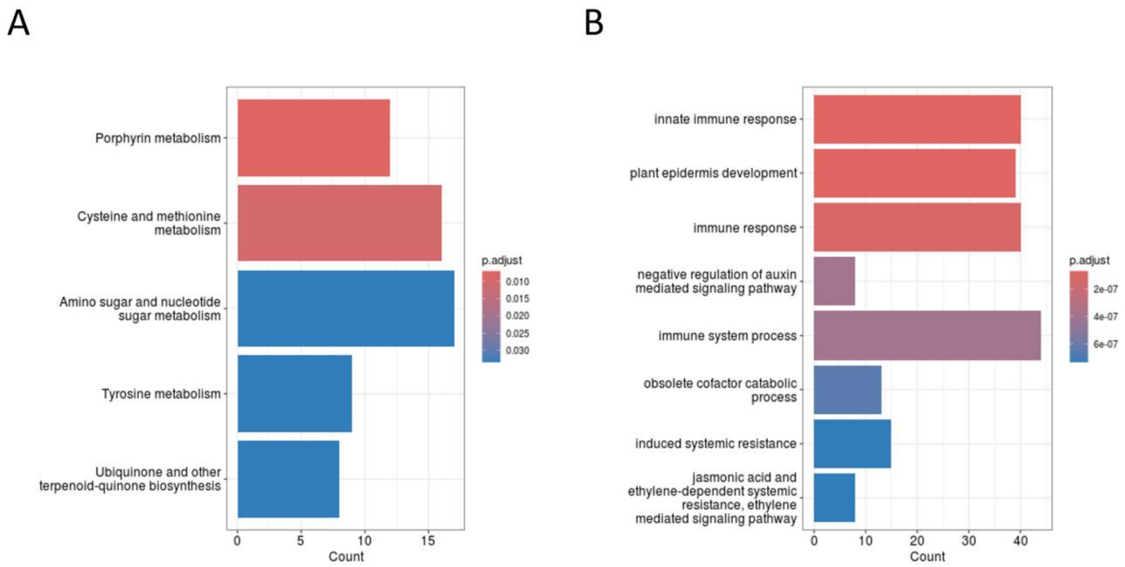


Supplementary Fig.19. GO and KEGG enrichment of unique and unclustered genes. A. GO enrichment bar plot. B. KEGG enrichment bar plot.

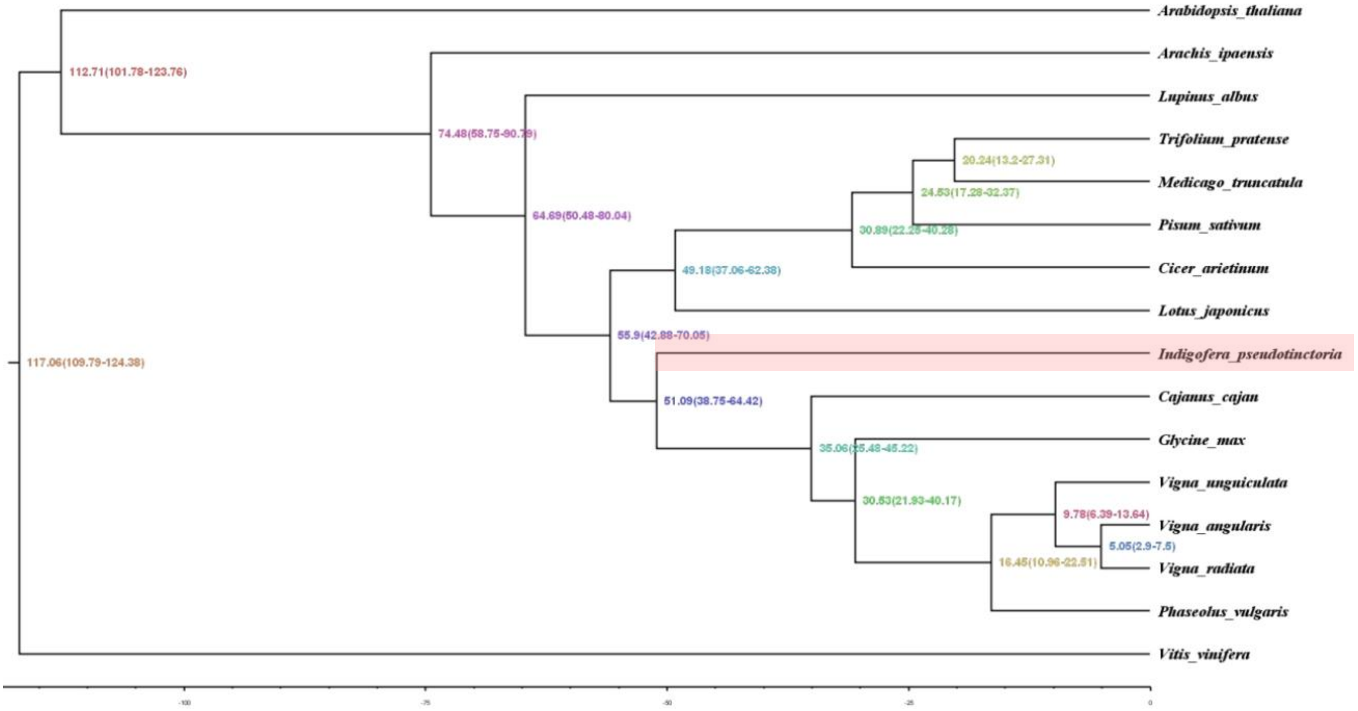


Supplementary Fig.20. Estimated species divergence times. Vitis vinifera was used as an outgroup. Numbers at each node represent estimated divergence times (in millions of years ago, Mya).

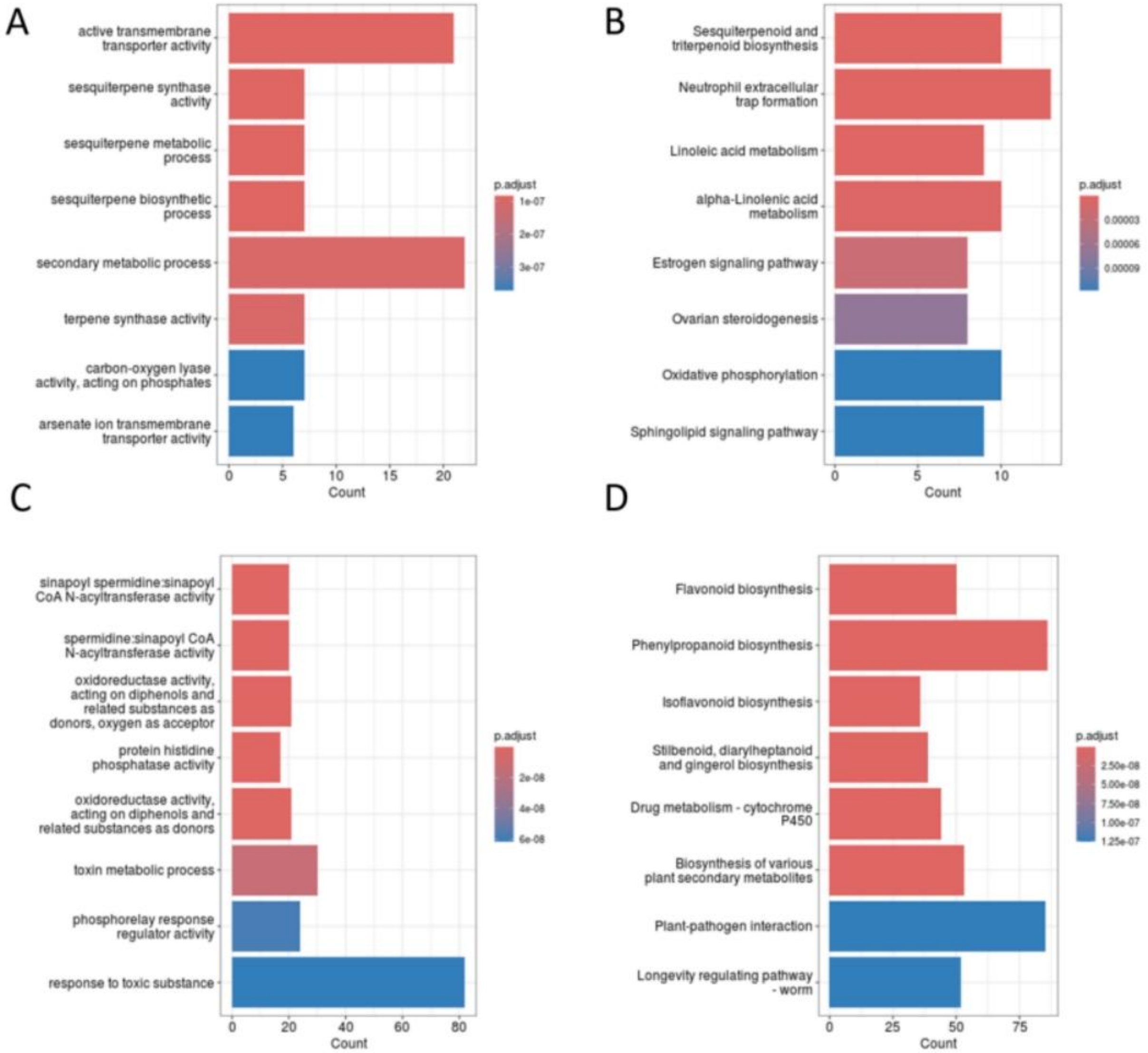


Supplementary Fig.21. GO and KEGG enrichment of significantly contracted and expanded gene families. A. GO classification bar plot of significantly contracted gene families. B. KEGG classification bar plot of significantly contracted gene families. C. GO classification bar plot of significantly expanded gene families. D. KEGG classification bar plot of significantly expanded gene families.

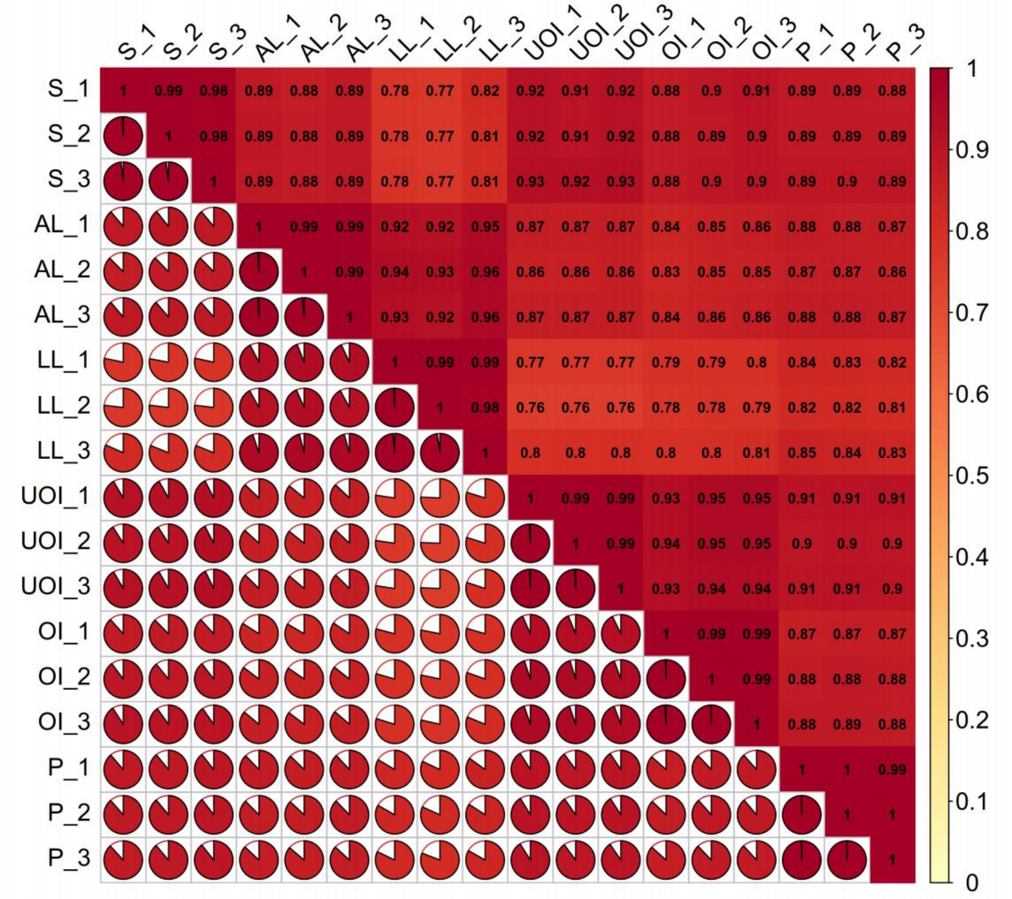

Supplementary Fig.22 The heatmap shows the correlation coefficients between gene expression levels across different tissue samples (S, AL, LL, UOI, OI, P) mapped to Hap1. The pie charts in each cell display the proportion of gene expression correlation between tissue pairs.

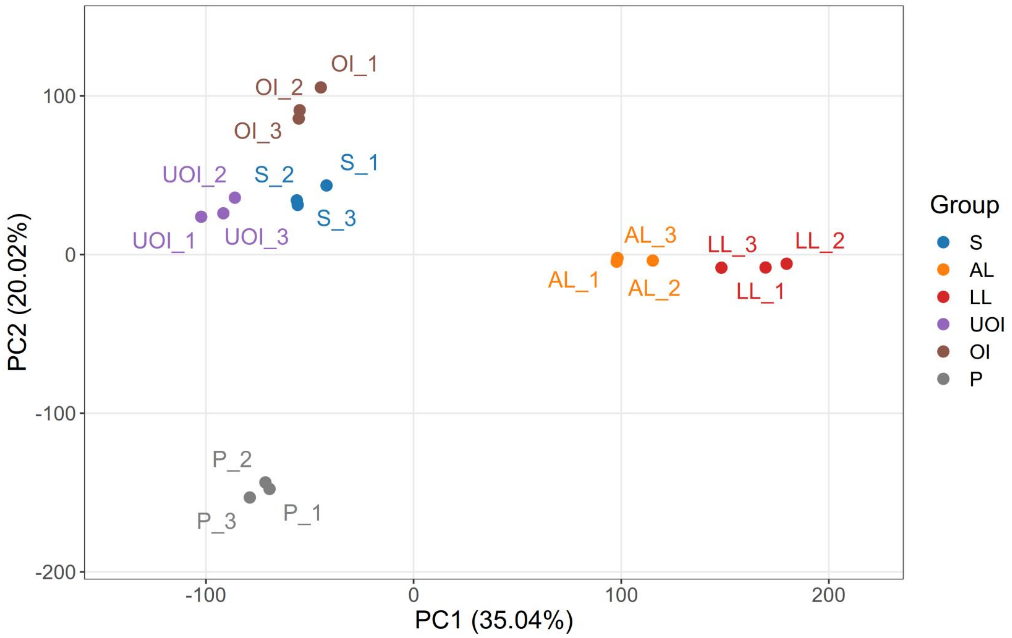


Supplementary Fig.23 Principal component analysis (PCA) of transcriptomic data mapped to Hap1 in *I. pseudotinctoria*. (S: blue, AL: orange, LL: red, UOI: purple, OI: brown, P: gray).

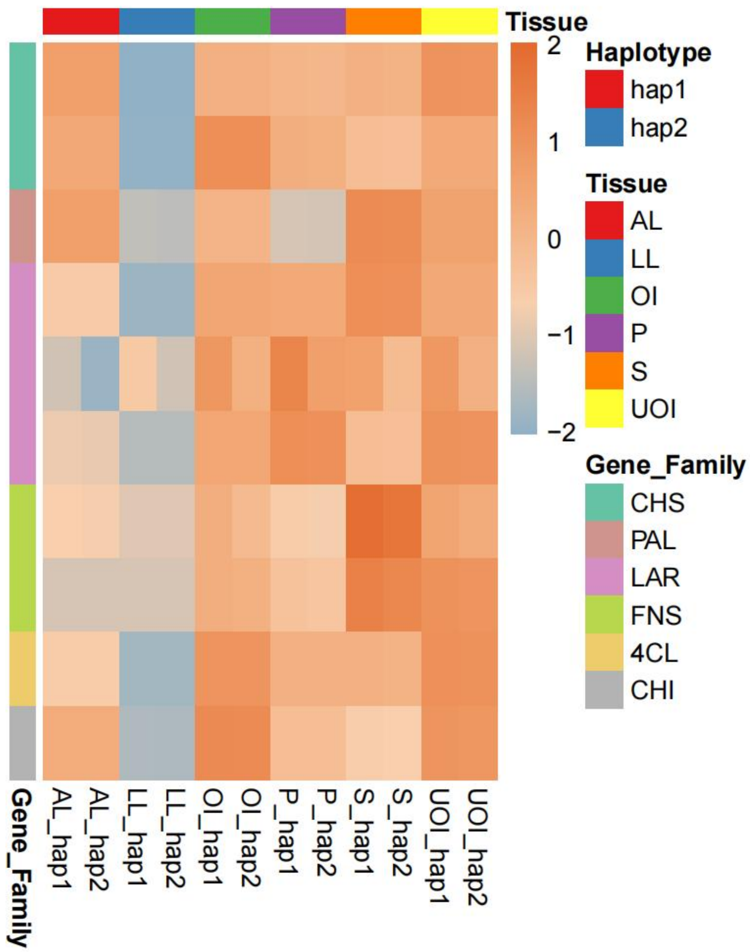


Supplementary Fig.25 Tissue-specific expression patterns of ten biallelic candidate genes shared by both haplotypes of *I. pseudotinctoria*.

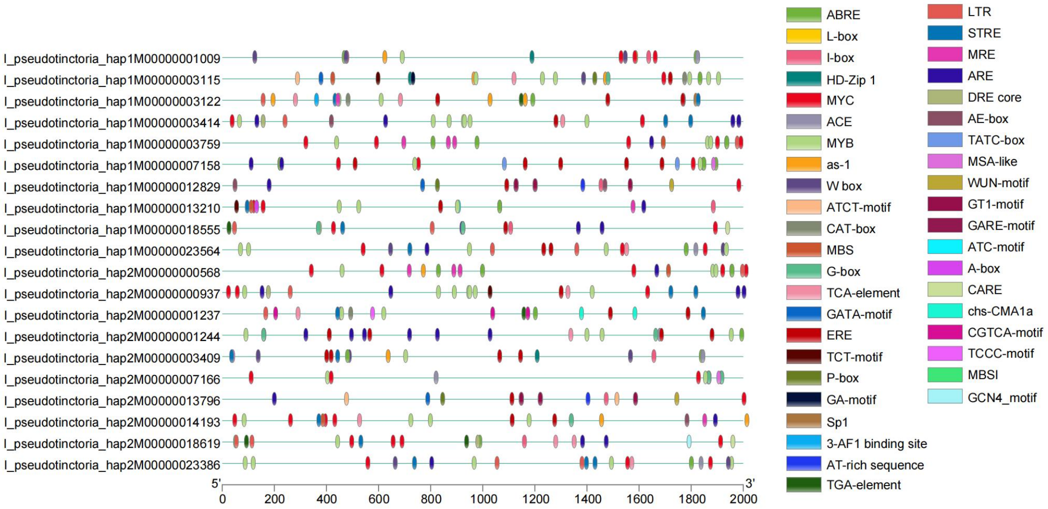


Supplementary Fig.26 Transcription factor binding sites within the 2 kb upstream region of candidate genes
